# scCancer2: data-driven in-depth annotations of the tumor microenvironment at single-level resolution

**DOI:** 10.1101/2023.08.22.554137

**Authors:** Zeyu Chen, Yuxin Miao, Zhiyuan Tan, Qifan Hu, Yanhong Wu, Wenbo Guo, Jin Gu

## Abstract

Single-cell RNA-seq (scRNA-seq) is a powerful technique for decoding the complex cellular compositions in the tumor microenvironment (TME). As previous studies have defined many meaningful cell subtypes in several tumor types, there is a great need to computationally transfer these labels to new datasets. Also, different studies used different approaches or criteria to define the cell subtypes for the same major cell lineages. The relationships between the cell subtypes defined in different studies should be carefully evaluated. In this updated package scCancer2, designed for integrative tumor scRNA-seq data analysis, we developed a supervised machine learning framework to annotate TME cells with annotated cell subtypes from 15 scRNA datasets with 594 samples in total. Based on the trained classifiers, we quantitatively constructed the similarity maps between the cell subtypes defined in different references by testing on all the 15 datasets. Secondly, to improve the identification of malignant cells, we designed a classifier by integrating large-scale pan-cancer TCGA bulk gene expression datasets and scRNA-seq datasets (10 cancer types, 159 samples, 652,160 cells). This classifier shows robust performances when no internal confidential reference cells are available. Thirdly, this package also integrated a module to process the seq-based spatial transcriptomic data and analyze the spatial features of TME. Software availability: http://lifeome.net/software/sccancer2/.

## 1 Introduction

The rapid development of single-cell RNA-sequencing (scRNA-seq) and spatial transcriptome (ST) technologies has promoted the accumulation of large-scale single-cell resolution omics data, enabling us to analyze the composition of tumor microenvironment (TME) more precisely and comprehensively.

Currently, researchers have defined many meaningful TME cell subtypes in various cancer types [1–3], which provides the opportunity to train supervised machine learning models and transfer these expert annotations to new datasets. Therefore, we upgraded our toolkit scCancer [4] to a new version (scCancer2) and improved the cell type annotation to more subtle subtypes. The updated version tended to preserve the labeling information or prior knowledge in the original references and assigned multi labels to the user’s data with a supervised machine learning framework. Compared with previous methods for cell type annotation [5–9], we focused on the scenarios of TME and provided a set of classifiers for annotating the cell types and subtypes defined in different references. As the accumulation of scRNA-seq based TME studies, there is a common need to examine the differences and similarities of the cell subtypes defined in different references. So, based on the trained classifiers, we established a cell subtype similarity map across multiple references by comparing the predicted labels on a large-scale tumor single cell collections (15 datasets, 594 samples and 1,213,469 cells; 5 training datasets for T cell subtypes, 3 for B cells, 6 for myeloid cells, 4 for endothelial cells and 3 for fibroblasts). The generated similarity map is a meaningful source to summarize and compare the abundant knowledge from different datasets.

At the single-cell level, in addition to cell subtype annotation, it is also crucial to identify the malignant cells in TME. Currently, the methods based on copy number variations (CNV) [4, 10, 11] have been applied to the malignancy annotation in various cancer types. However, when no internal confidential reference cells are available, CNV-based methods frequently get unstable results. To be compatible with these special scenarios, we developed an additional data-driven method. Firstly, a reference dataset combining scRNA-seq dataset (652,160 cells in 159 samples) and bulk RNA-seq data (7,012 TCGA samples) was established across multiple cancer types and then an XGBoost [12] based classifier was trained to identify malignant cells with high generalization ability. Besides, the classifier achieved higher computational efficiency and lower memory burden, which is suitable for processing large-scale datasets. To demonstrate the robustness of this malignant cell identification module in scCancer2, we categorized the test samples into four groups, including tumor samples with the bimodal or unimodal distribution of malignancy score, normal samples, and organoid samples. Extensive tests proved that it can be a great supplement to CNV-based methods.

Finally, the spatial dimension is also highly significant in characterizing the TME. To systematically dissect the TME spatial features, we constructed an automated spatial transcriptome analysis module, which includes three analytical perspectives, including spatial interaction, spatial heterogeneity, and spatial structure. By applying it to 60 samples from various cancer types, we demonstrated its good performance.

Overall, taking advantage of the accumulated scRNA-seq data of cancer clinical samples, the in-depth expert annotations for TME in each study, and the new spatial information brought by the ST technology, scCancer2 improved its functionalities for automatically dissecting the complex TME features. The performances of scCancer2 have been extensively tested on a large amount of data of various cancer types.

## 2 Results

### 2.1 Overview of scCancer2

We implemented three new modules in our R toolkit scCancer2 (**Figure 1**). The first module: scCancer2 automatically performed cell subtype classification with a series of built-in models and output multi-label annotations. To visualize the relationship of labels quantitatively, a similarity map of cell subtypes from different atlases was calculated for every major cell type. The second module: scCancer2 identified malignant cells based on bulk RNA-seq and scRNA-seq with the XGBoost. Our method can be directly applied to various types of samples without relying on the inference of CNVs. The third module: scCancer2 performed systematical analyses of spatial transcriptomics data for cancer research, mainly consisting of basic analysis and TME spatial analysis (interaction, heterogeneity, and structure). The information on packages and methods newly integrated into scCancer2 were summarized in **Table S1**.

**Figure 1:**
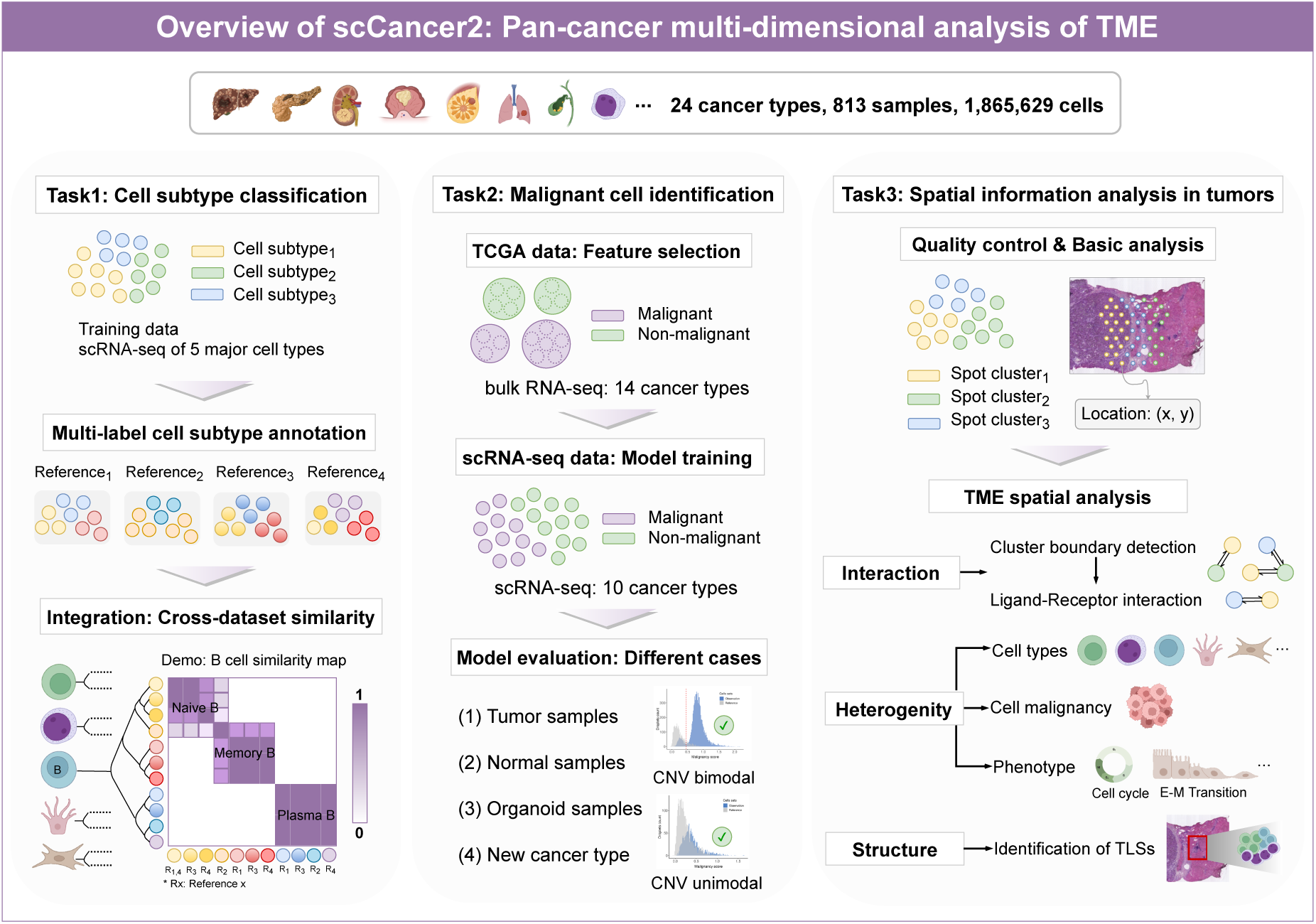
The overview of scCancer2.

### 2.2 scCancer2 upgrades tumor microenvironment cell type annotation to the subtype level

Advances in scRNA-seq enabled researchers to provide accurate depictions of TME. The cell annotation of recently published datasets was usually precise to the cell subtype level, which can be served as reference datasets for cell subtype classification. We collected and preprocessed a total of 15 published datasets across 17 human cancer types and 3 sequencing technologies (10X Genomics, Smart-seq2, InDrop). There are 594 samples with 1,213,469 cells in total. (**Table 1**)

**Table 1:** The list of scRNA-seq datasets used in cell subtype annotation.

| Accession | Organ | No. of cells | No. of samples | Cell type | Technology |
| --- | --- | --- | --- | --- | --- |
| GSE98638 [1] | Liver | 5,063 | 6 | T cell | Smart-Seq2 |
| GSE99254 [2] | Lung | 12,346 | 14 | T cell | Smart-Seq2 |
| GSE108989 [3] | Colorectal | 11,138 | 12 | T cell | Smart-Seq2 |
| GSE178341 [13] | Colorectal | 371,223 | 181 | all | 10X |
| GSE176078 [14] | Breast | 100,064 | 26 | all | 10X |
| GSE154763 [15] | Pan-cancer | 65,083 | 48 | myeloid cell | 10X |
| GSE127465 [16] | Lung | 54,773 | 40 | immune cell | inDrop |
| GSE150430 [17] | Nasopharyngeal | 48,584 | 16 | immune cell | 10X |
| GSE146771 [18] | Colorectal | 54,285 | 20 | all | Smart-Seq2, 10X |
| GSE140228 [19] | Liver | 73,286 | 41 | immune cell | Smart-Seq2, 10X |
| GSE103322 [20] | Head and Neck | 5,902 | 18 | all | Smart-Seq2 |
| GSE132465 [21] | Colorectal | 65,362 | 33 | all | 10X |
| GSE131907 [22] | Lung | 208,506 | 58 | all | 10X |
| E-MTAB-6149 [23] | Lung | 40,250 | 19 | stromal cell | 10X |
| GSE156728 [24] | Pan-cancer | 97,604 | 62 | T cell | 10X |

<sup>1</sup> In the column "Cell type", "all" represents 5 major cell types: immune cells (T cells, B cells, and myeloid cells) and stromal cells (endothelial cells and fibroblasts). Epithelial cells are excluded.
<sup>2</sup> In the column "No. of cells", the numbers represent the total number of cells in the datasets. For model training and validation, we usually selected specific cell types (a subset of the original dataset).
<sup>3</sup> GSE156728 was treated as the query dataset. The patient samples were not used for model training.

Here, we implemented a rapid, supervised annotation framework integrating prior information. The training step for every single dataset mainly includes the following steps. Firstly, select cells by the expression of marker genes to construct high-quality training sets [7]. Secondly, select genes according to the statistic entropy [25]. Thirdly, train a multinomial model for each cell subtype and estimate the expression probability of the selected genes by maximum likelihood estimation (**Section 4.1**). Then, for each major cell type, we trained a series of subtype annotation models based on multiple reference datasets (**Section 4.1**).

By applying to 594 samples in **Table 1**, we proved that scCancer2 had great generalization ability on TME datasets of 5 major cell types (**Figure 2A**), including T cells, B cells, myeloid cells, endothelial cells and fibroblasts. Moreover, we compared the framework with other benchmarks including scPred [6], one-class logistic regression (OCLR) [26], multinomial, and Scibet [25] on 6 datasets containing 5 major cell types and 3 sequencing technologies. Considering the prediction accuracy, computational efficiency, and model complexity, the results in **Figure 2B** further indicate the reliability of our framework. Taking 43,817 immune cells in colorectal cancer (CRC) dataset GSE146771 [18] as an example, cell types were annotated hierarchically. We annotated major immune cell types through OCLR in scCancer [4]. Then, for each major cell type, we assigned multiple sets of cell subtype labels based on multiple trained models in scCancer2 and visualized them respectively (**Figure 2C**). Correspondence between the suffix number of labels and reference datasets can be found in **Table S2**.

**Figure 2:**
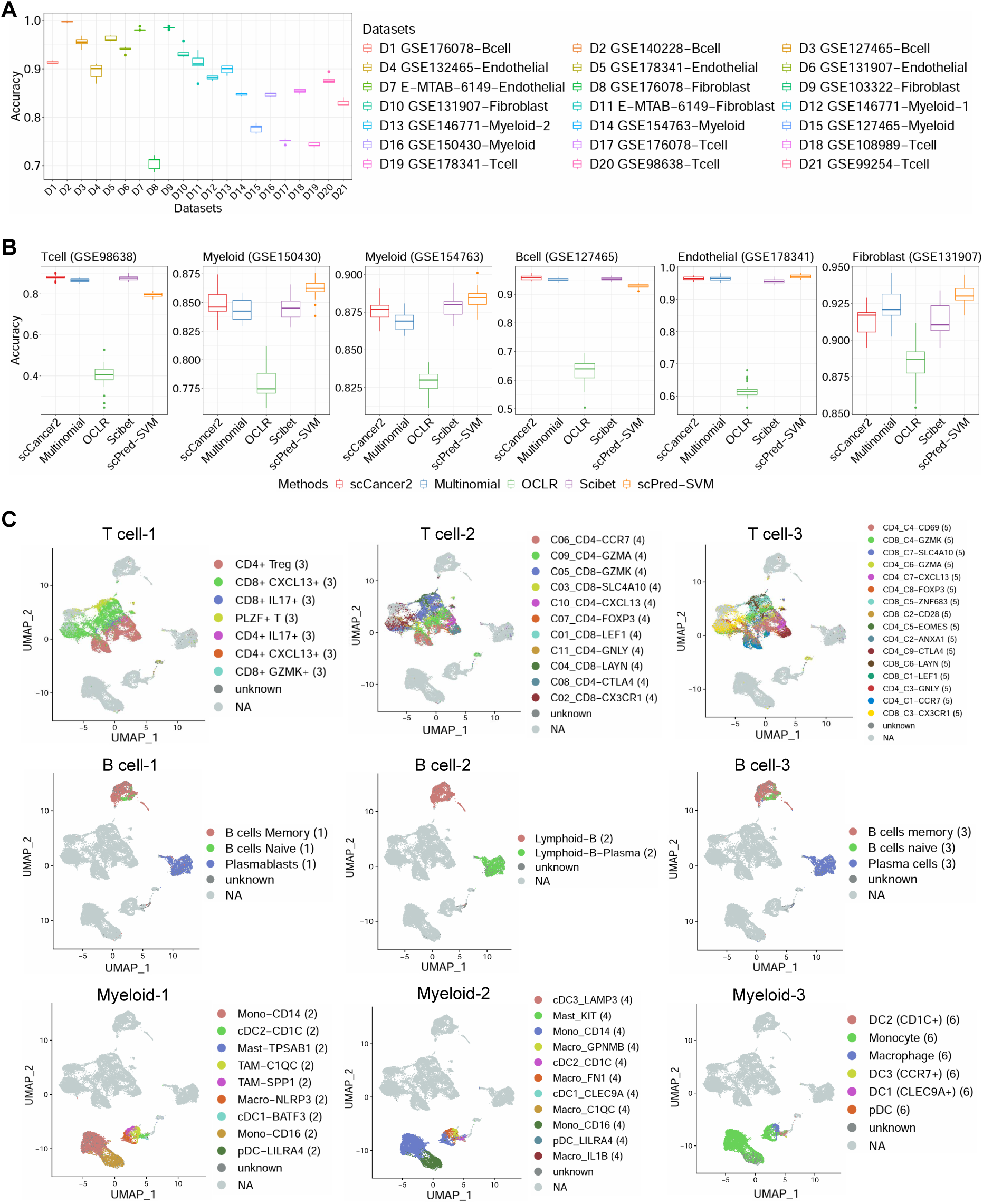
scCancer2 annotates cell subtypes of tumor microenvironment. **(A)** Performance evaluation of scCancer2 on cell subtype annotation task. The results were obtained by 5-fold cross-validation. **(B)** Comparison of different machine learning methods on cell subtype annotation task. The results were obtained by 5-fold cross-validation. **(C)** Multi label annotation results on CRC example (immune cells). Rough cell labels (T cell, myeloid cell, B cell) were assigned to CRC example through OCLR in scCancer. The dataset was then divided by major cell types. Cell subtype annotation was achieved by scCancer2, forming a multi-model hierarchical cell type annotation. In each subfigure, the numbers after the label represent the literature serial number corresponding to the major cell type.

In summary, we fully preserved the original annotation from different cell atlases and transferred them to new datasets with a supervised machine-learning framework.

### 2.3 scCancer2 quantitatively evaluates the similarity of cell subtypes across datasets

The automatic annotation system worked based on a series of built-in models trained through the above pipeline. scCancer2 built a multi-label annotation structure where each cell was assigned multiple possible labels, retaining the original annotation in cell atlases. However, the cellular composition of different cancer types varied greatly. There are significant differences in expert annotations at the subtype level among different datasets. Therefore, depicting the relationship between cell subtypes defined in different studies is urgently in need.

We carried out a method to integrate the cell annotations from different experts. We annotated cell subtypes across datasets with those built-in models respectively and each label was assigned to several cells. Then, we extracted “label-barcode” sets from the results and derived the Jacobian similarity. Finally, the Jacobian similarity matrix was visualized by heatmap, hierarchical clustering, and multidimensional scaling (**Section 4.1**). By observing the aggregation of labels in 2-dimensional space, we can understand the complicated relationship between different cell subtypes.

Take CRC dataset [18] as an example, cell annotation was conducted based on multiple “pre-trained” models, and a cross-dataset label relationship of B cells was calculated, as shown in **Figure 3A**. It was observed that the aggregation pattern of B cell labels was consistent with expectations: 2 naive B cell labels, 3 plasma cell labels, and 2 memory B cell labels showed a high degree of similarity. In particular, the “Lymphoid-B” in HCC was expected to be the union of näıve B cells and memory B cells in NSCLC and BRCA. Respectively, its similarity with the naive B cells was 0.10 and 0.17, and that with the memory B cells was 0.72 and 0.67 (**Figure 3A**). The results indicate the inclusion relationship between these groups of labels, which is consistent with the biological meaning of the labels. In the right panel of **Figure 3A**, subtypes of B cells can be clearly divided into 3 categories based on spatial distance. The “Lymphoid-B” label was located between the naive B category and the memory B category and closer to the latter.

**Figure 3:**
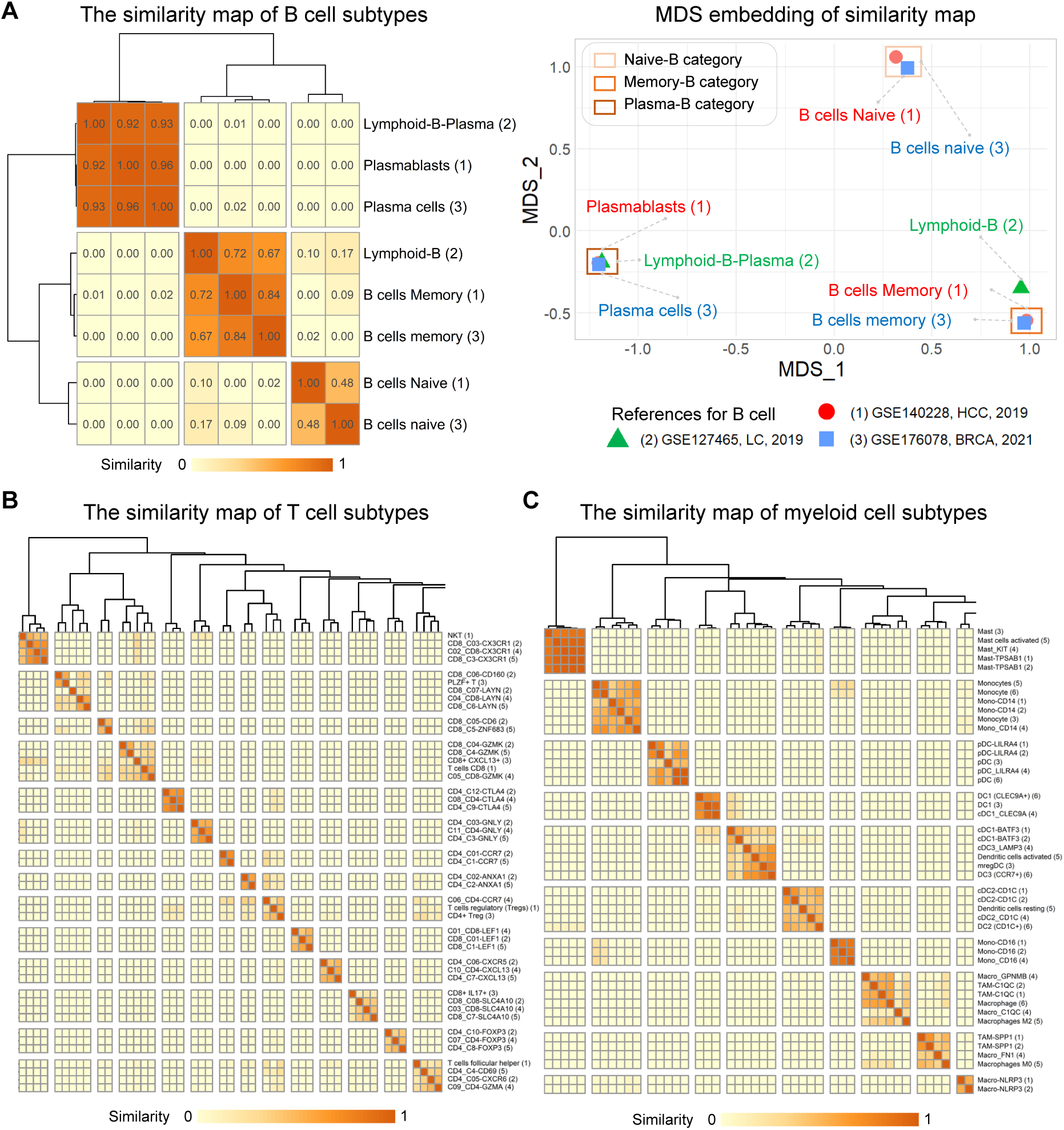
Generation of cross-dataset cell subtype similarity by scCancer2. **(A)** The cross-dataset similarity map of lymphoid B cell subtypes on CRC-example. The left panel is the heatmap of the similarity matrix. The right panel is the multidimensional scaling plot of the matrix, demonstrating the similarity of different labels. Color represents the source of subtype labels. **(B)** The cross-dataset similarity map of lymphoid T cell subtypes on CRC-example. **(C)** The cross-dataset similarity map of myeloid cell subtypes on CRC-example. * In each subfigure, the numbers after the label represent the literature serial number corresponding to the major cell lineage. The horizontal and vertical coordinates of the heatmap are the same. Each point in the matrix represents the Jaccord similarity between the horizontal axis label and the vertical axis label, and the diagonal element is 1. **(B)** and **(C)** are part of the similarity map (See **Figure S2**-**Figure S6** and **Table S2** for full version and references).

Similarly, the similarity maps of T cells and myeloid cells were generated. We have taken a portion of the complete similarity graph in **Figure 3B** and **Figure 3C**. Firstly, cell sub-types from different studies of the same group of researchers were often named in a similar pattern. They were almost all clustered into the same category in the similarity map. For example, “CD4 C12-CTLA4”, “C08 CD4-CTLA4”, and “CD4 C9-CTLA4” in **Figure 3B**. Except for identical labels mentioned above, cell subtypes named by different researchers with similar functions also showed high similarity in the heatmap. For example, “Mast”, “Mast cells activated”, “Mast KIT”, and “MAST-TPSAB1” in **Figure 3C**. Finally, we integrated the original annotations of the pan-cancer query dataset [24] into the similarity graph. Observing the results in **Figure S1**, we found that almost all newly assigned labels aggregated in the heatmap had a corresponding original annotation. The annotation of scCancer2 had a good consistency with the original annotations. These results indicate that scCancer2 successfully depicted the cross-dataset relationship of complicated immune cell subtypes with similar functions (See **Figure S2**-**Figure S6** for all results and the corresponding studies).

In summary, we quantitatively depicted the connections between cell subtypes from different cell atlases with the label similarity graph. We extracted and summarized the abundant information in the multi-label annotation structure. It can be an important reference and literature indexing tool.

### 2.4 scCancer2 identifies malignant cells across multiple cancer types without internal references

It is an important and challenging problem to determine whether a cell is malignant, which is of key significance not only for subsequent analysis of tumor heterogeneity and microenvironmental characteristics but also for the study of the mechanism of tumor occurrence and development. scCancer [4] scored cell malignancy based on inferCNV [10] and compared the distribution of malignancy scores with normal references. Malignancy labels were assigned based on a bimodality observation of scores. However, copy number variation is not always necessary for cell malignancy. When there is no internal confidential reference cell, the CNV-based method becomes unreliable. For example, when the proportion of tumor and normal cells in a sample is seriously imbalanced, the malignancy scores follow a unimodal distribution.

To extract the transcriptome characteristics of malignant cells, we designed a pipeline to identify malignant cells from scRNA-seq data and TCGA data with the machine learning method XGBoost, as a supplement to the CNV-based method. The pipeline included reference dataset annotation based on the distribution of malignancy score, feature selection based on cancer-specific features extracted from the TCGA dataset, model training, and performance evaluation (**Section 4.2**). We collected a total of 652,160 cells in 159 samples (**Table S3**) and constructed a high-quality single-cell reference dataset with 466,468 cells and 2,756 genes. To ensure the reliability of the classifier, it is vital to balance the proportion of normal and malignant cells. Moreover, multiple cancer types were required in the training and evaluation sets for better pan-cancer generalization ability. Consequently, we divided the training and validation sets by sample sources instead of random sampling (**Table S3**).

Then, ElasticNet, XGBoost, and Neural Network were applied respectively to identify malignant cells in scRNA-seq data. The results show that XGBoost achieved better performance compared with ElasticNet and Neural Network (**Figure 4A**). In terms of efficiency, the XG-Boost significantly improves computational efficiency and reduces memory usage compared with the CNV-based method, especially when processing integrated large-scale datasets [18, 27]. By observing the significant genes selected by the XGBoost during the training process, we qualitatively judged the consistency between biological prior knowledge and the data-driven method. In particular, we noted that cancer-related genes including *METTL1*, *ALKBH2*, *PSPH*, *PT-PRC*, and *TRIP13* contributed the most to the identification, which indicated the biological interpretability of our model (**Figure 4B**).

**Figure 4:**
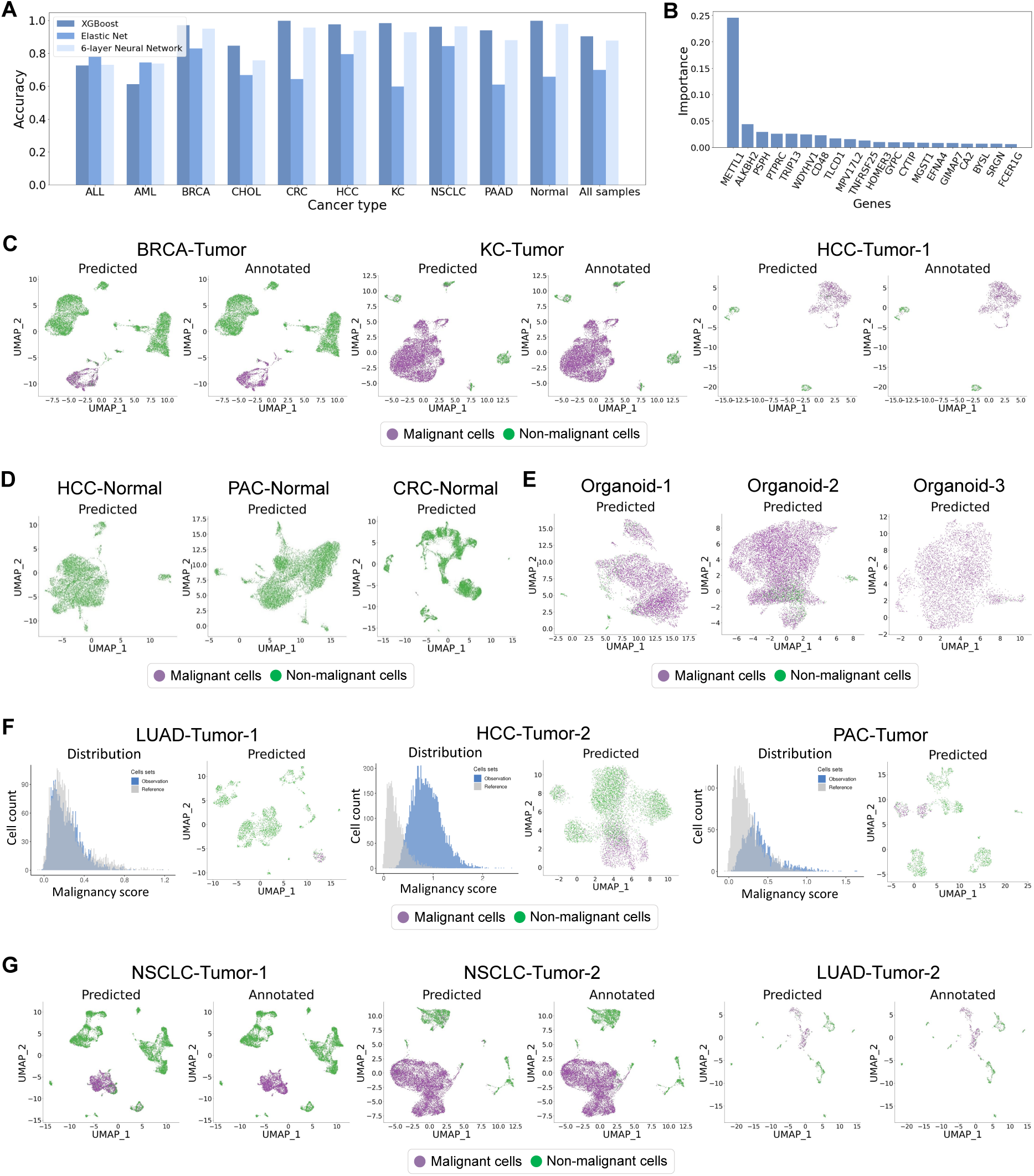
The results of malignant cell identification by scCancer2. **(A)** Performance comparison of different machine learning methods across multiple cancer types on malignant cell identification task. **(B)** Important genes for cell malignancy. The histogram shows the feature importance ranking calculated by the XGBoost model during the training process. **(C)** Case 1. Test scCancer2 on tumor samples with bimodal distribution of malignancy score. For each subfigure (cancer type), the left panel shows the annotation results of scCancer2 (XGBoost), while the right panel shows the annotation results of scCancer (CNV-based method). **(D)** Case 2. Test scCancer2 on normal samples. The UMAP-plot shows the annotation results of scCancer2 on 3 samples without malignant cells. **(E)** Case 3. Test scCancer2 on organoid samples. The UMAP-plot shows the annotation results of scCancer2 on 3 *in vitro* cholangiocarcinoma organoid samples. **(F)** Case 4. Test scCancer2 on tumor samples with unimodal distribution of malignancy score. For each subfigure, the left panel shows the distribution of malignancy score, while the right panel shows the annotation results of scCancer2. **(G)** Case 5. Test scCancer2 on new cancer types. The subfigures compare the prediction results and ground truth on 3 lung cancer samples. See **Table S3** for the correspondence between the original name of the datasets and their naming in figures.

We applied the model to several test samples. When the malignancy scores of tumor samples have a significant bimodal distribution, the annotation results of scCancer2 (XGBoost) on malignant cells are highly consistent with the annotation results of scCancer [4] (CNV-based method) (**Figure 4C** and **Figure S7**). When the proportion of malignant cells and normal cells is imbalanced (normal samples and cholangiocarcinoma organoid samples), scCancer2 can accurately identify normal cells (**Figure 4D** and **Figure S8**) and malignant cells (**Figure 4E** and **Figure S8**), which can be a supplement to the CNV-based method. Through the annotation of tumor organoid samples, we can evaluate the quality of *in vitro* culture. Furthermore, the CNV-based method is unreliable for a minority of solid tumor samples. When the malignancy score of the sample follows a unimodal distribution, scCancer2 can also identify malignant cells (**Figure 4F**). We found that the identified malignant clusters were highly consistent with epithelial cells predicted by scCancer [4] in solid tumor samples (**Figure S9**). Lastly, scCancer2 can effectively predict cancer types that do not exist in the training set. In the experiment, we moved the lung cancer (LC) samples from the training set into the validation set. Then, we trained a new model and directly predicted the malignant cells in LC samples. We found that our model still performed well on these samples (**Figure 4G**).

In summary, the results indicate that scCancer2 effectively extracted the transcriptome characteristics of malignant cells in TME, obtaining great generalization ability across cancer types. It is a great supplement and improvement to the CNV-based method.

### 2.5 scCancer2 analyzes tumor microenvironment from a spatial perspective

Sequencing-based spatial transcriptomics has helped researchers to achieve impressive results in exploring the spatial heterogeneity of malignant tumors and TME [28]. scCancer2 provides a highly automated module for tumor spatial transcriptome analysis, which mainly consists of two parts. The first part focuses on quality control (QC), statistical analysis of spots and genes, and basic downstream analysis. The second part analyzes TME from three different angles: spatial interaction, spatial heterogeneity, and spatial structure.

We first utilized the spatial information to perform morphology QC to filter small isolated tissue areas, which often do not contribute to the analysis results. Then, we visualized unique molecular identifier (UMI) counts and detected gene numbers to reflect characteristics of the tissue to a certain extent and filtered the spots with extremely low UMI counts or gene numbers. We provided statistical results of UMI counts on 60 tumor samples from 9 cancer types (**Figure S10** and **Table S5**). Furthermore, we performed statistical analysis for gene expression proportion especially for mitochondrial genes and ribosomal genes (**Figure S11**). After QC, we performed basic downstream analyses including dimension reduction, clustering, and differential expression analysis based on Seurat [29].

TME spatial analysis module includes ligand-receptor interaction, tumor heterogeneity, and spatial structure detection. The interaction of different regions is crucial to understand tumor behaviors, such as growth, progression, drug response, and therapeutic effect. Therefore, scCancer2 identified the boundary spots between clusters automatically and defined the ligand-receptor interaction strength between two clusters as the mean of the average expression of the ligand and receptor pairs from the CellPhoneDB dataset [30]. As seen in **Figure 5A**, scCancer2 marked the boundary between the fibrous capsule and the tumor. The results of ligand-receptor interaction indicate that the extracellular matrix receptor interaction (collagen-related ligand-receptor pairs) was strong at the boundary. Besides, several disease-related ligand-receptor pairs such as CD74-APP/COPA/MIF and CXCL12-ACKR3/DPP4 were also found at the boundary [31–33]. Moreover, marker genes of common cell types [34] were used to roughly estimate the cell distribution and cell composition in each spot. We defined the cell type evaluation scores as the average expression of marker genes (**Figure S12**). To further explore tumor heterogeneity, we considered 14 phenotypes from CancerSEA [35]. For example, we found differences between regions with malignant cell enrichment and others. In HCC-1L, areas with dense distribution of malignant cells had higher cell cycling scores (**Figure 5B**).

**Figure 5:**
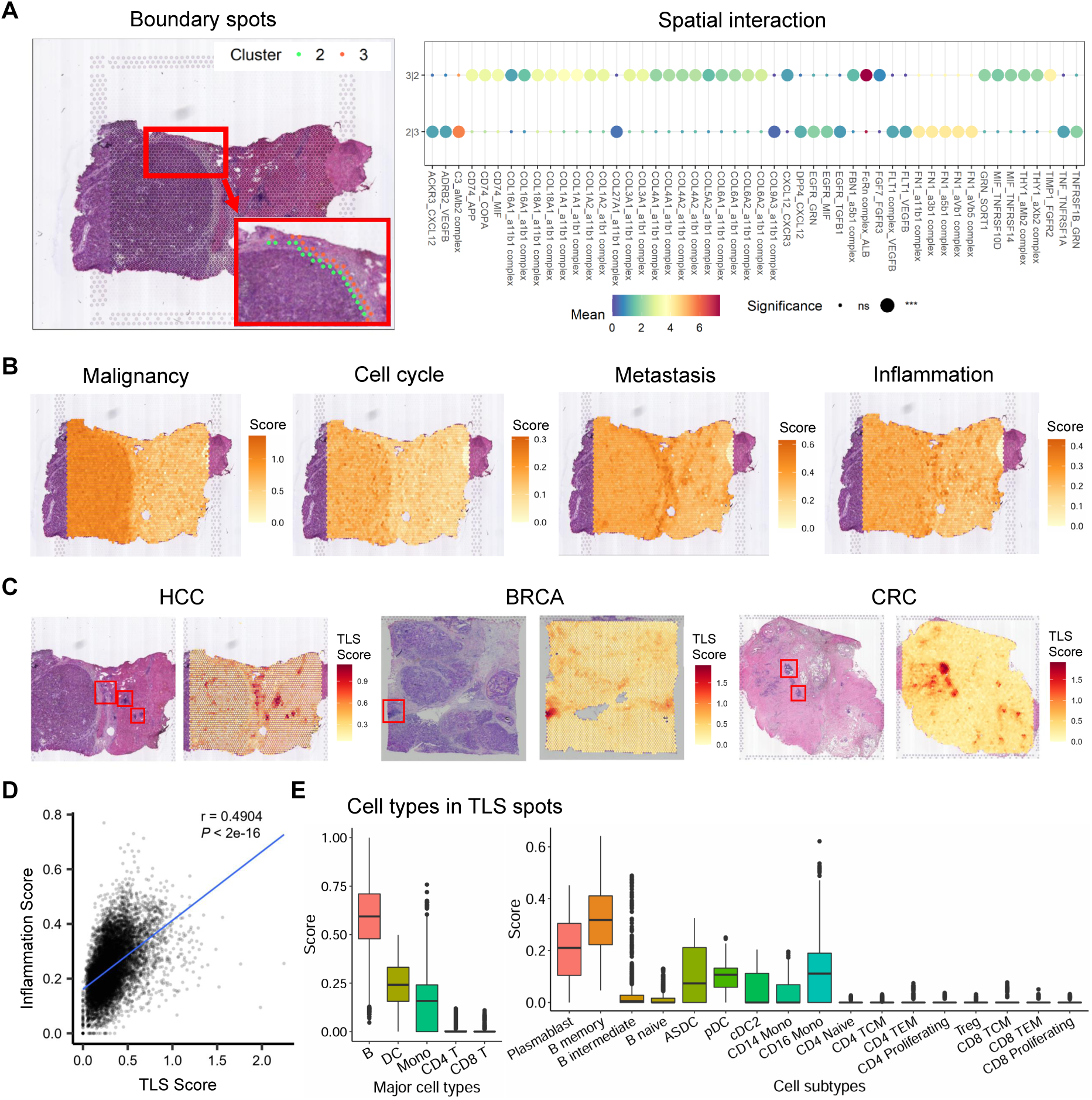
The results of TME spatial analysis by scCancer2 on HCC examples. **(A)** Boundary spots of neighboring clusters (left panel) and the ligand-receptor interaction strength between two clusters (right panel) in HCC-1L. **(B)** Tumor phenotype heterogeneity analysis by scCancer2. Cell malignancy, cell cycle, metastasis, and inflammation were scored in HCC-1L based on marker genes. **(C)** Tertiary lymphoid structures identified by scCancer2 and their corresponding H&E images. HCC: “HCC-1L” in Wu et al. [34]; BRCA: “V1 Breast Cancer Block A Section 2” in 10x Genomics; CRC: “ST-colon3” in Wu et al. [40]. **(D)** Correlation analysis between spatial structure (TLSs) and spatial characteristic (inflammation) on samples across various cancer types. **(E)** Major cell lineages and cell subtypes enriched in TLSs. The boxplot contained all major and minor cell types which predicted to be positive in the TLSs.

Tertiary lymphoid structures (TLSs) in the TME correlate with clinical outcomes and arise in the context of chronic inflammation and represent prognostic biomarkers and cancer treatment targets [36]. scCancer2 detected TLS regions using the average expression of TLS-50 signature [34] and compared them with corresponding H&E images of tumor samples (**Figure 5C** and **Figure S13**). It achieved good performance on various cancer types, including HCC, BRCA, CRC, etc. Furthermore, we also found that there was a significant correlation between spatial structures and transcriptome characteristics. The detected distribution of TLSs was correlated with the distribution of inflammatory features in 60 tumor samples across 9 cancer types (**Figure 5D**).

Finally, we extended the module of cell subtype annotation to the spatial level by quantitatively evaluating the cellular composition of TLSs. We detected TLSs from tumor samples and transferred the annotations of cell subtypes from the reference dataset [37] to all TLS spots (**Section 4.3**). As shown in the first panel of **Figure 5E**, we detected 4 enriched immune cell types in the spots of TLSs, including B cells, dendritic cells, monocytes, and T cells, which is consistent with previous studies [38, 39]. The cell types not presented were all 0-score, such as NK cells. At the cell subtype level, plasmablasts, memory B cells, and dendritic cells are the most abundant. As shown in the second panel of **Figure 5E**, common subtypes of T cells can also be detected.

In summary, we implemented a new module to analyze TME with sequencing-based spatial transcriptomics. This module is highly systematic and automated. It enables us to explore the spatial characteristics of TME from multiple perspectives comprehensively.

## 3 Discussion

By leveraging machine learning and integrating multiple analysis modules, we provided a valuable tool scCancer2 for researchers to gain a more comprehensive understanding of the TME at the single-cell level. Our analysis consists of three different angles. Firstly, we trained a series of machine-learning models on scRNA-seq data for cell subtype annotation and quantitatively evaluated the similarity of labels originating from different datasets. Secondly, we constructed a reference dataset based on bulk RNA-seq and scRNA-seq data and identified the malignant cells with the XGBoost model. Thirdly, we integrated spatial transcriptome analysis modules for a multi-dimensional view of TME.

For the cell subtype annotation module, we have tested classic machine learning methods (for example, SVM and XGBoost) and specialized algorithms for scRNA-seq on massive data. The evaluation of classification algorithms requires consideration of multiple factors: classification accuracy, computational efficiency, model complexity, and biological interpretability. In terms of accuracy within a single dataset, scCancer2 is close to SVM in scPred [6]. However, scCancer2 can generate lightweight models and rapidly assign multiple sets of labels to new input data. Meanwhile, we found that XGBoost achieved better performance than other algorithms on classification tasks (**Figure S14**), but its generalization ability was worse than scCancer2 on cross-dataset annotation, making it hard to discover the relationship of labels originating from different datasets (**Figure S15** and **Figure S16**).

The results of cross-dataset annotation indicate the generalization ability of scCancer2 on immune cells and endothelial cells. Cell subtypes with similar functions showed high similarity in the heatmap (**Figure S2**-**Figure S5**). scCancer2 provides a convenient access for users to not only extract specific cell subtypes from the dataset but also index similar subtypes and relevant references. The results in fibroblasts are relatively ambiguous (**Figure S6**) due to the heterogeneity of fibroblasts [41, 42]. We hope to propose reliable indicators for quantitatively evaluating the cross-dataset performance of algorithms based on similarity maps.

For the malignant cell identification module, machine learning-based methods in scCancer2 can be a great supplement to the CNV-based method. Firstly, scCancer2 extracts the transcriptome characteristics of malignant cells and performs well when the CNV-based method fails to work. Secondly, when directly processing large-scale published datasets instead of individual samples, the time cost and memory burden of the CNV-based method are high, while the trained model in scCancer2 can rapidly identify malignant cells in large-scale datasets.

For the spatial transcriptome analysis module, exploring tumor heterogeneity from a spatial perspective is significant. As we have already analyzed the composition of TLSs at the cell subtype level, we hope to benchmark more deconvolution and mapping strategies for the integration of scRNA-seq and spatial transcriptomics data [43]. Transcriptome characteristics extracted from bulk RNA-seq and scRNA-seq data such as malignancy features can also be utilized for spatial heterogeneity analysis.

In the future, we will pay more attention to the scalability of scCancer2. In recent studies, there are several newly discovered or rare cell subtypes [44, 45]. Detecting them in the existing samples is an important issue. We can treat these cell subtypes as the positively labeled queries and solve it with novelty detection methods such as one-class models [46]. Moreover, adding new subtypes to existing label networks rapidly (without cross-dataset annotation) is challenging. Co-modeling the expression matrix and the label similarity matrix as a label network by graph models can not only visualize the relationship more clearly, but also provide access for new nodes by graph embedding and link prediction.

## 4 Methods

### 4.1 TME cell subtype annotation

#### Cell subtype annotation method within a single dataset

Firstly, we randomly divided the dataset into training and validation sets in a 4:1 ratio (5-fold cross-validation). Given a training expression matrix *X* of cell subtype labels *L* and the marker genes of every subtype *G_prior_* (or differentially expressed genes of clusters), we first selected representative cells *C* to construct core training data according to the marker score [7]. Then, the Entropy test [25] or Highly Regional Genes [47] was used for feature selection. The genes used for training were the union of input markers and genes selected from the expression matrix.

Therefore, we obtained an expression matrix *X_C,G_* for model training. We trained a multinomial model in which every parameter represents the expression probability *P* of a corresponding gene. For the validation set, we also evaluated the expression of marker genes. We labeled a cell as “unknown” when it did not express any marker gene in the prior information, which means that for every possible cell subtype, the ‘Aggregate Marker Score’ function output a low marker score. Then, we assigned cell subtypes to other unlabeled cells with maximum likelihood estimation. *P* was weighted by the marker score mentioned above.

We evaluated the performance of our model through 5-fold cross-validation on 15 published datasets, using the mean and variance of classification accuracy as indicators. During the training process, we obtained 21 cell subtype templates across 5 major cell types in total. In addition, we compared our pipeline with multinomial in Scibet [25], OCLR [26] in scCancer [4], SVM in scPred [6], and XGBoost [12] on 6 datasets with 5 major cell types and 3 sequencing technologies.

#### Multi-label annotation structure

For the newly input dataset *D*, we first annotated the cell types with scCancer [4] and split the dataset based on major cell types. For every subset *D_sub_*, we traversed all corresponding cell subtype templates and annotated cell subtypes with the above pipeline iteratively. The output is summarized in **Table 2** below.

**Table 2:**
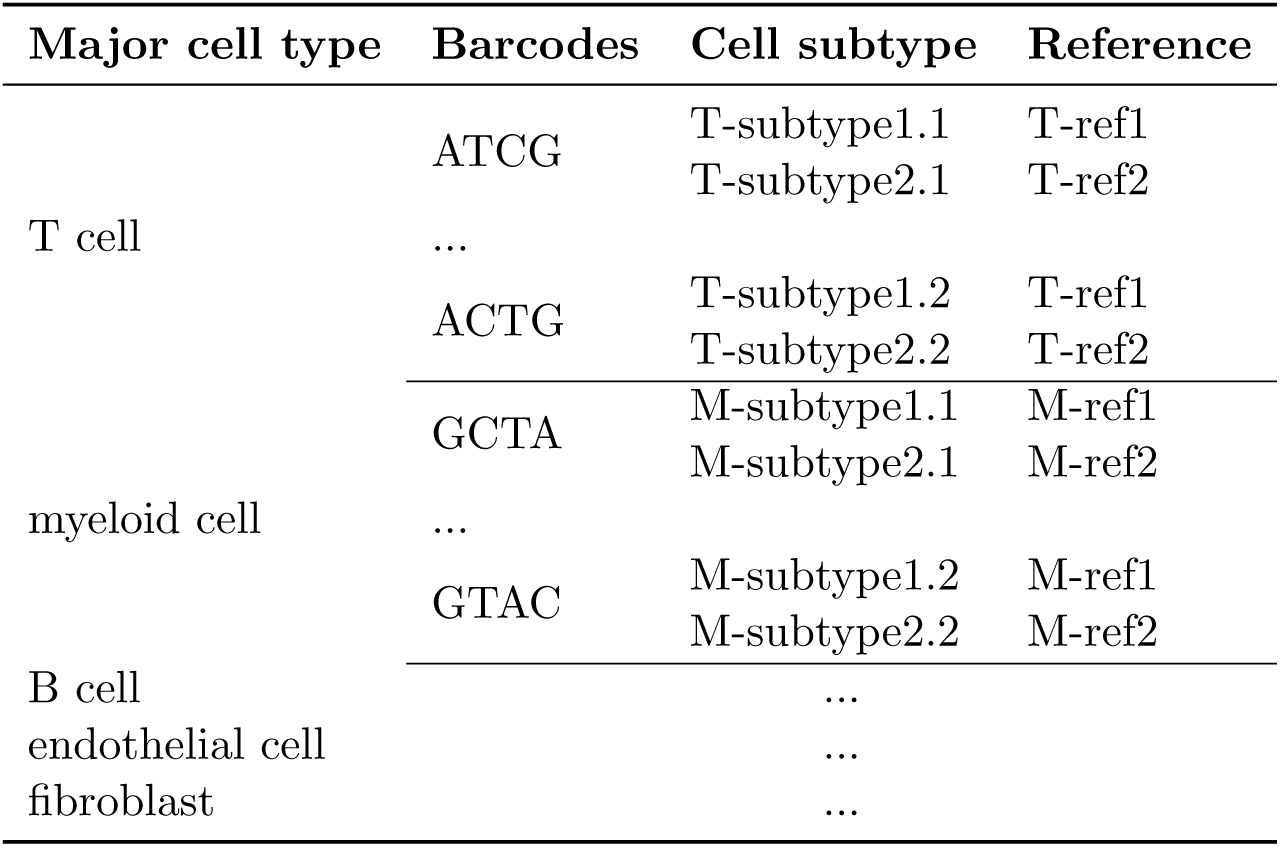
Multi-label annotation for input TME dataset.

#### Similarity map of different subtype labels

Defining a set *C_ij_*as cells assigned to the *j^th^*subtype in the *i_th_*group of labels, namely, the *i^th^*dataset, and the similarity matrix as *S*, the similarity of every 2 labels was calculated as follows:

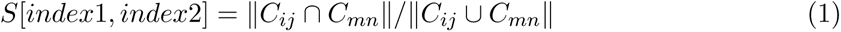

Where the index is the position of a subtype after flattening all the subtype labels into a vector, and the dimension of matrix *S* is the length of the label vector, namely, the total number of subtype labels. Cross-annotation process within the training set is summarized in **Algorithm 1**. Only the requirement changes for the new test dataset: 1 test dataset, *N* annotation models, and *N* groups of labels.

##### Algorithm 1

Multi-model and multi-dataset cross annotation

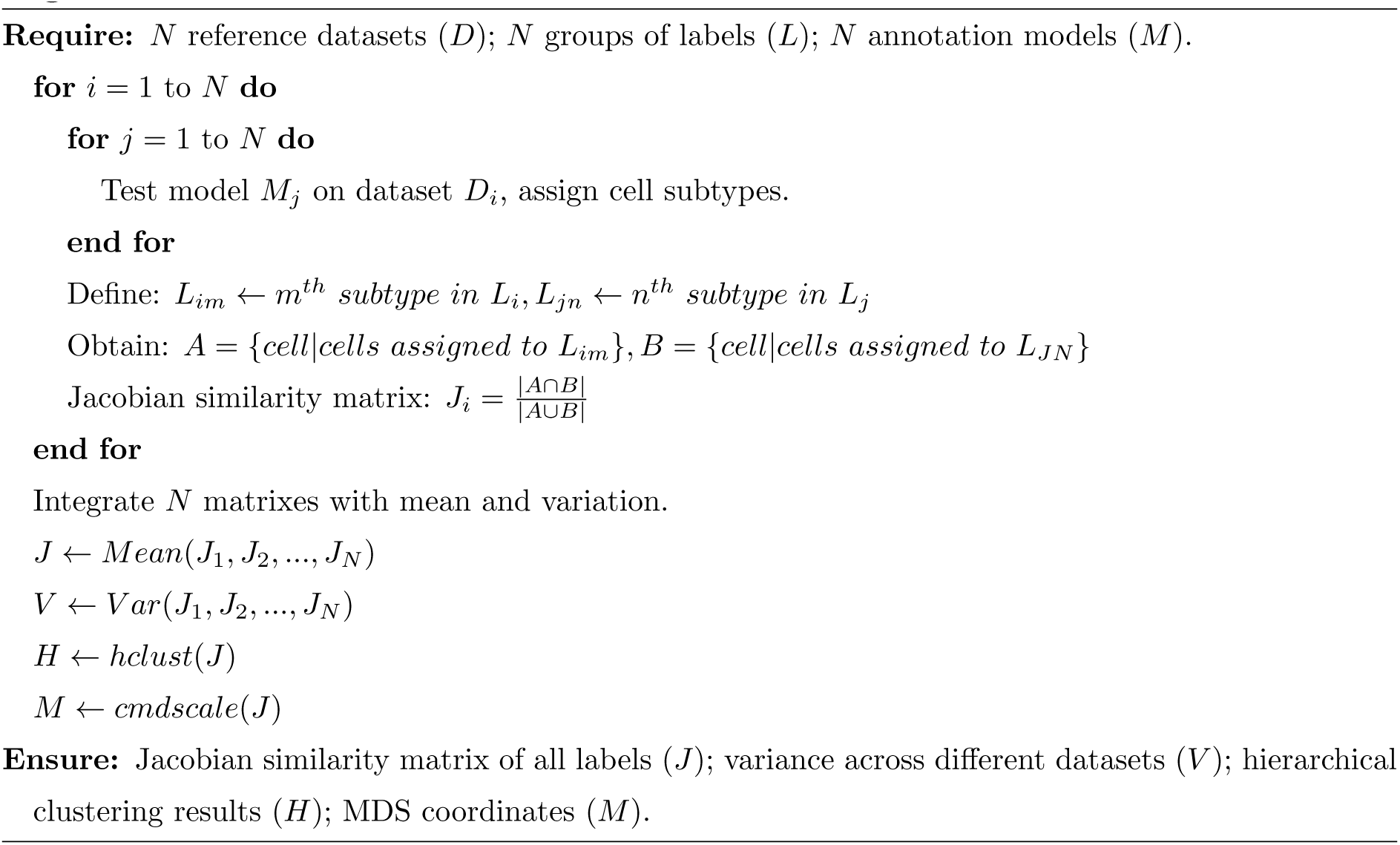

### 4.2 Malignant cell identification

#### Data collection, preprocessing, and annotation

We first collected tumor samples sequenced by 10X Genomics with no previous malignant annotation. We calculated the copy number variation score of tumor samples through the inferCNV [10] and plot the distribution of malignancy scores. We retained samples with an obvious bimodal distribution.

Although inferCNV has been proven to be reliable when the malignancy score follows a bimodal distribution, we found that almost all malignant cells are epithelial in the labeled results. To prevent the model from regarding the features of epithelial cells as malignant, we collected normal samples with more normal epithelial cells. Additionally, we downloaded and integrated published datasets from the TISCH2 database [48] as a supplement. (**Table S3**) To integrate samples from different sources, we preprocessed the data through scanpy [49] and concatenated the expression matrixes based on common genes.

#### Feature selection combining bulk RNA-seq and scRNA-seq

We first collected bulk RNA-seq data of 14 cancer types from TCGA. Then we pre-processed and integrated the samples based on common genes. We calculated differentially expressed genes between malignant samples and normal samples across 14 cancer types using the edgeR [50] package. From these groups of differentially expressed genes, we identified high-frequency genes, named as *G_bulkDE_*.

As for scRNA-seq data, we selected 2,000 genes with the highest expression variance as *G_scHV_*. The features selected from scRNA-seq data *G_scRNA_* for model training were the union of *G_bulkDE_* and *G_scHV_*.

#### Model training and performance evaluation

We divided the training set and validation set by sample sources (**Table S3**) and trained an XGBoost model with a binary logistic objective function. We analyzed the performance of scCancer2 under various scenarios: tumor samples with bimodal distribution of malignancy scores, tumor samples with unimodal distribution of malignancy scores, normal samples, and organoid samples. In addition, we evaluated the generalization ability of scCancer2 by changing the cancer types in the training set.

### 4.3 Spatial transcriptome analysis

#### Data collection

The test datasets for spatial module in scCancer2 were obtained from previous studies and 10X Genomics. We collected 60 samples across 9 cancer types in total. (**Table S5**)

#### Quality control

We first utilized spatial information to perform morphological quality control on spots. In some samples, isolated tissue areas may appear in the section, which often does not contribute to the analysis results, and even bring errors to clustering, and differential gene analysis. Therefore, we removed these small regions through connected domain processing.

For 10X Visium data, we obtained the 6 nearest neighbors of each spot and constructed connected domains. By default, regions with connected domain area less than 3 are removed. The remaining area will be used for the following steps. Then, we visualized the detected gene numbers and nUMI in the tissue image and filtered the spots with extremely low gene numbers or nUMI. In addition to QC on spots, we also performed QC on genes. First, we filtered genes that expressed in less than 3 cells. Then we represented the expressions of mitochondrial genes and ribosomal genes, and it is suggested that researchers carefully consider the effects of these genes on the final results.

#### Basic downstream analysis

After quality control steps, we performed basic downstream analyses based on Seurat, including normalization, highly variable genes identification, dimension reduction (PCA/tSNE/UMAP), clustering, and differential expression analysis. Then we represented them with some redesigned visualization functions.

#### Gene expression program analysis

Non-negative matrix factorization (NMF) [51] can identify gene expression programs unsupervisedly. Each column of the left matrix obtained after NMF decomposition can be regarded as the relevant gene of a program extracted, and each column of the right matrix can be regarded as the strength of each program expressed by the tissue. In scCancer2, we applied NMF decomposition to the normalized expression matrix to reveal expression programs. And the intensities of each expression program are shown in figures.

#### Spatial interaction analysis

The interaction of different regions is crucial to understand some tumor behaviors, such as growth, progression, drug response, and therapeutic effect. Here, we referred to CellPhoneDB to characterize ligand-receptor interactions extent over the boundaries of clusters. Firstly, scCancer2 identified the boundary spots between clusters automatically. Then, we extracted expressions of the approximately 1,400 known ligand-receptor pairs from the CellPhoneDB dataset, and defined the ligand-receptor interaction score between two clusters as the mean of the average expression of the ligand and receptor genes in the boundary spots of each cluster, respectively. To identify significant interactions, we randomly disrupted the labels of clusters 1,000 times, calculated the strength of each interaction pair in each cluster under the new labels each time, formed a null distribution of 1,000 sampling points for each interaction pair, and filtered weak gene pairs according to their strengths got p-values of each interaction pair. Finally, we visualized these significant interactions between clusters by a scatter plot.

#### Phenotypic heterogeneity analysis

Intra-tumor heterogeneity is widespread in cancer samples and highly associated with poor prognosis of cancer. The spatial transcriptomics technologies provide an avenue to explore heterogeneity at the space level. In scCancer2, we mainly considered 14 characters from CancerSEA [35]. We integrated two methods to compare phenotypic scores in different spatial regions. The first is the ‘average’ method, which calculates the average expression of signature genes of each phenotype. The other one is the ‘AddModeuleScore’ function in Seurat.

#### Copy number variation analysis

Copy number variation is the potential to identify malignant tumor cells, which was widely used in the research of various cancer types. Here, we integrated the algorithm of R packages inferCNV and CopyKAT [11] to estimate CNVs of each spot. InferCNV was to calculate the moving average of expression values across each chromosome and then compared them with normal reference data to estimate CNVs. Considering the impact of dropout, we used spots’ neighbor information of PCA space to smooth CNV values. CopyKAT combined a Bayesian approach with hierarchical clustering to calculate genomic copy number profiles of single cells and defined clonal substructure. This method only required a gene expression matrix of UMI counts and then outputted the copy number profiles and predicted labels of cells. If users have reference data, we recommend using inferCNV because it is faster. If not, we recommend using CopyKAT to calculate CNVs automatically.

#### Detection of tertiary lymphoid structures

We collected the TLS-50 signature from Wu et al. [34] and located the TLS spots based on the distribution of TLS-50 scores (average gene expression). The threshold for filtering TLSs was manually set based on pathological images and gene expression scores. Next, we combined the tumor samples from Wu et al. [31], Wu et al. [34], and Wu et al. [40] with common genes. Then, we analyzed the cellular composition of the detected TLS spots with ‘FindTransferAnchors’ and ‘TransferData’ in Seurat[37]. We chose a multi-modal PBMC dataset in Hao et al. [37] as the reference and map the cell annotations to the combined TLS spots. To save memory, we extracted 50% of the original dataset by cell subtypes with the stratified sampling method.

## Code availability

The R package is available online. Package for the updated version of scCancer: http://lifeome.net/software/sccancer2/.

## Author Contributions

JG conceived the project. JG and WBG supervised the research. ZYC and YXM conducted the research. ZYC implemented the modules of cell subtype and cell malignancy. YXM implemented the module of spatial transcriptome analysis. ZYT contributed to the investigation and data processing of malignant cell identification. QFH and YHW contributed to the testing of scCancer2. ZYC and YXM wrote the original draft. JG, WBG, QFH and YHW contributed to the writing of the manuscript. All authors read and approved the final manuscript.

## Acknowledgements

This work was funded by the National Key Research and Development Program of China (Nos. 2020YFA0712403, 2021YFF1200901), the National Natural Science Foundation of China (Nos. 62133006, 92268104), the Tsinghua University Initiative Scientific Research Program (No. 20221080076) and the China Postdoctoral Science Foundation (No. 2022M721839).

## Supplementary Information

### Supplemental Figures

#### Similarity map of major cell subtypes on query datasets

**Figure S1:**
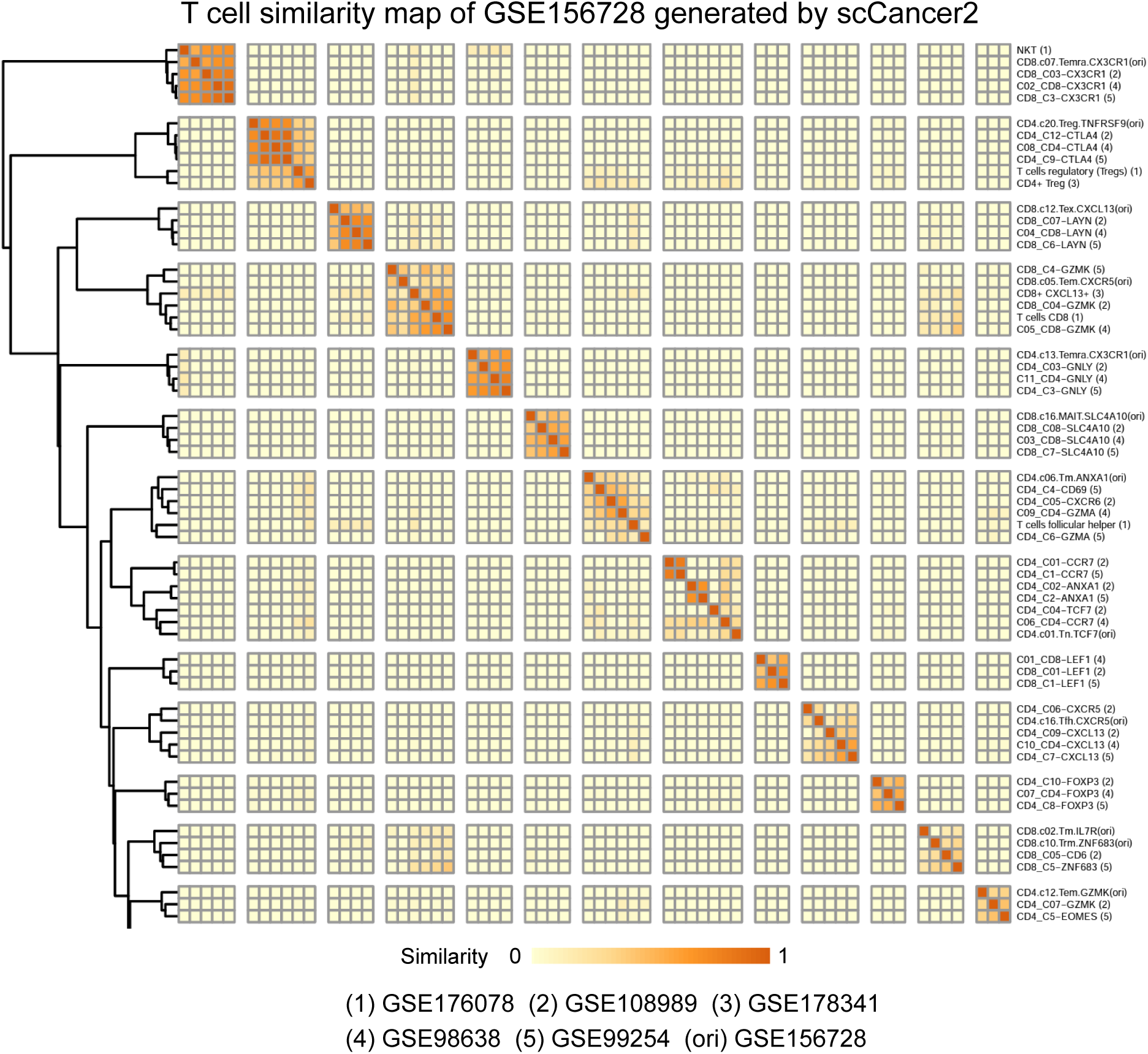
Similarity map of T cell subtypes on GSE156728. The original annotation of T cells were used for comparison.

#### Integrated similarity map of major cell subtypes on multiple training datasets generated by scCancer2

**Figure S2:**
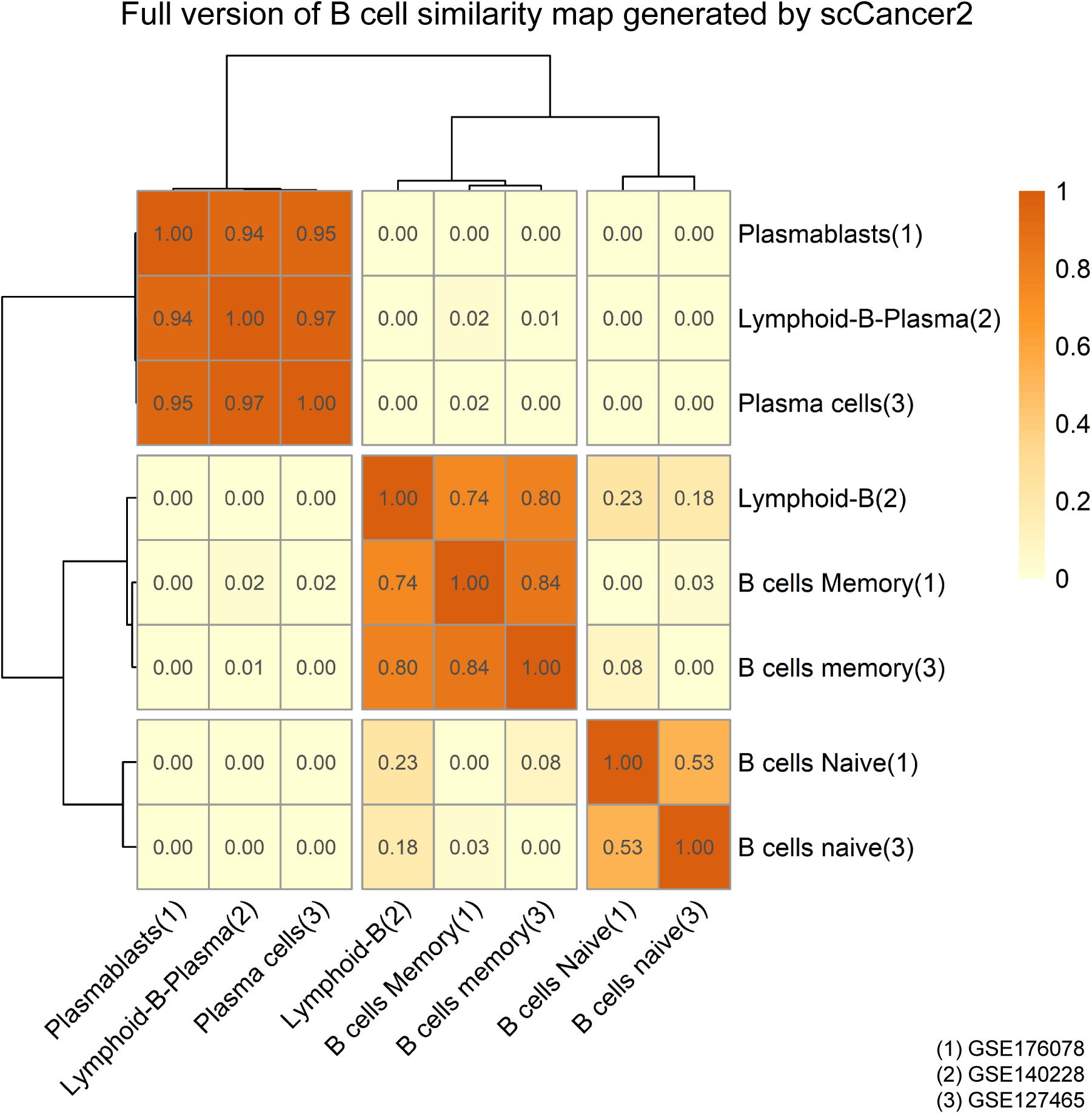
Similarity map of B cell subtypes generated from cross-dataset annotation (3 training sets).

**Figure S3:**
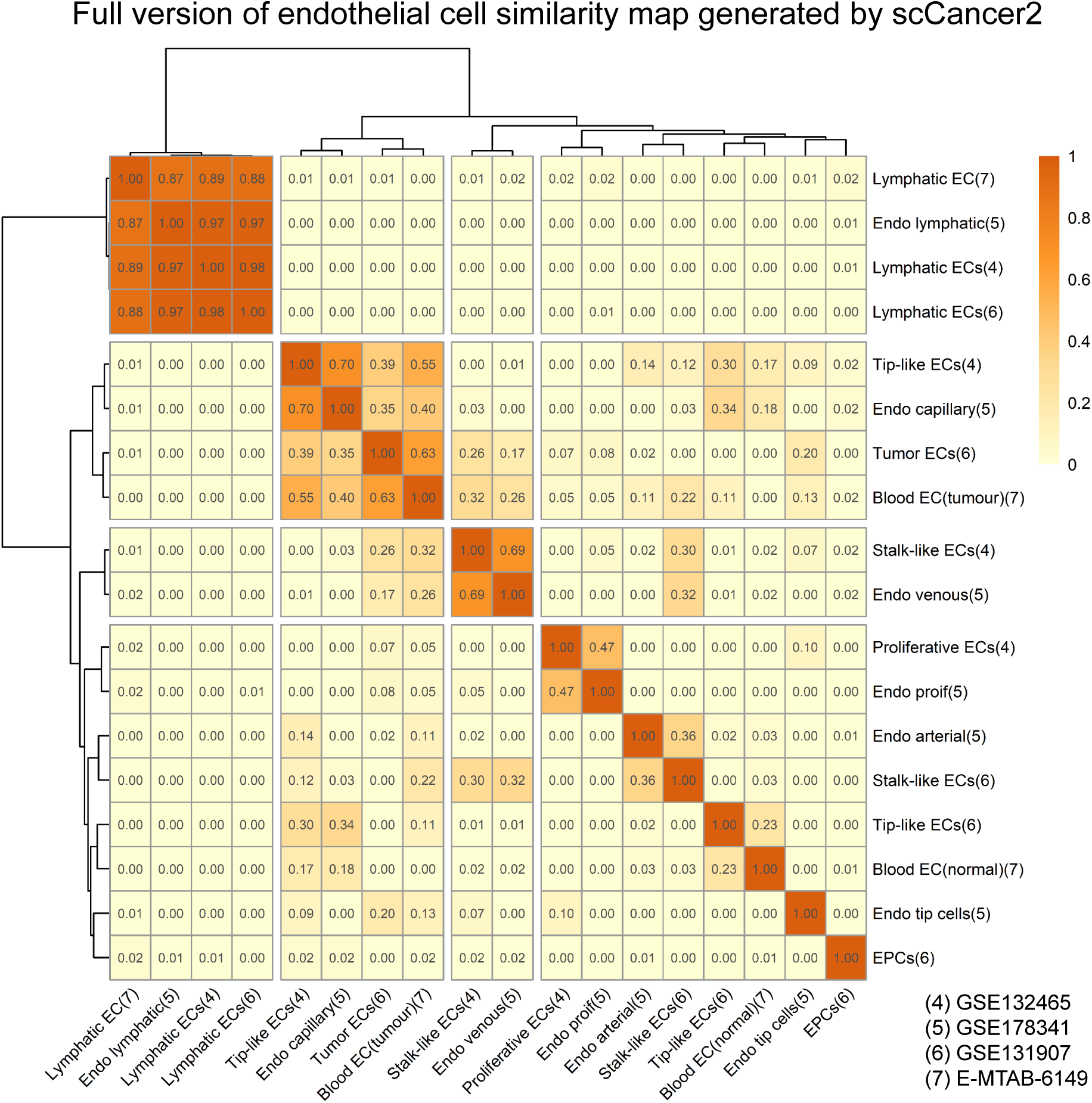
Similarity map of endothelial subtypes generated from cross-dataset annotation (4 training sets).

**Figure S4:**
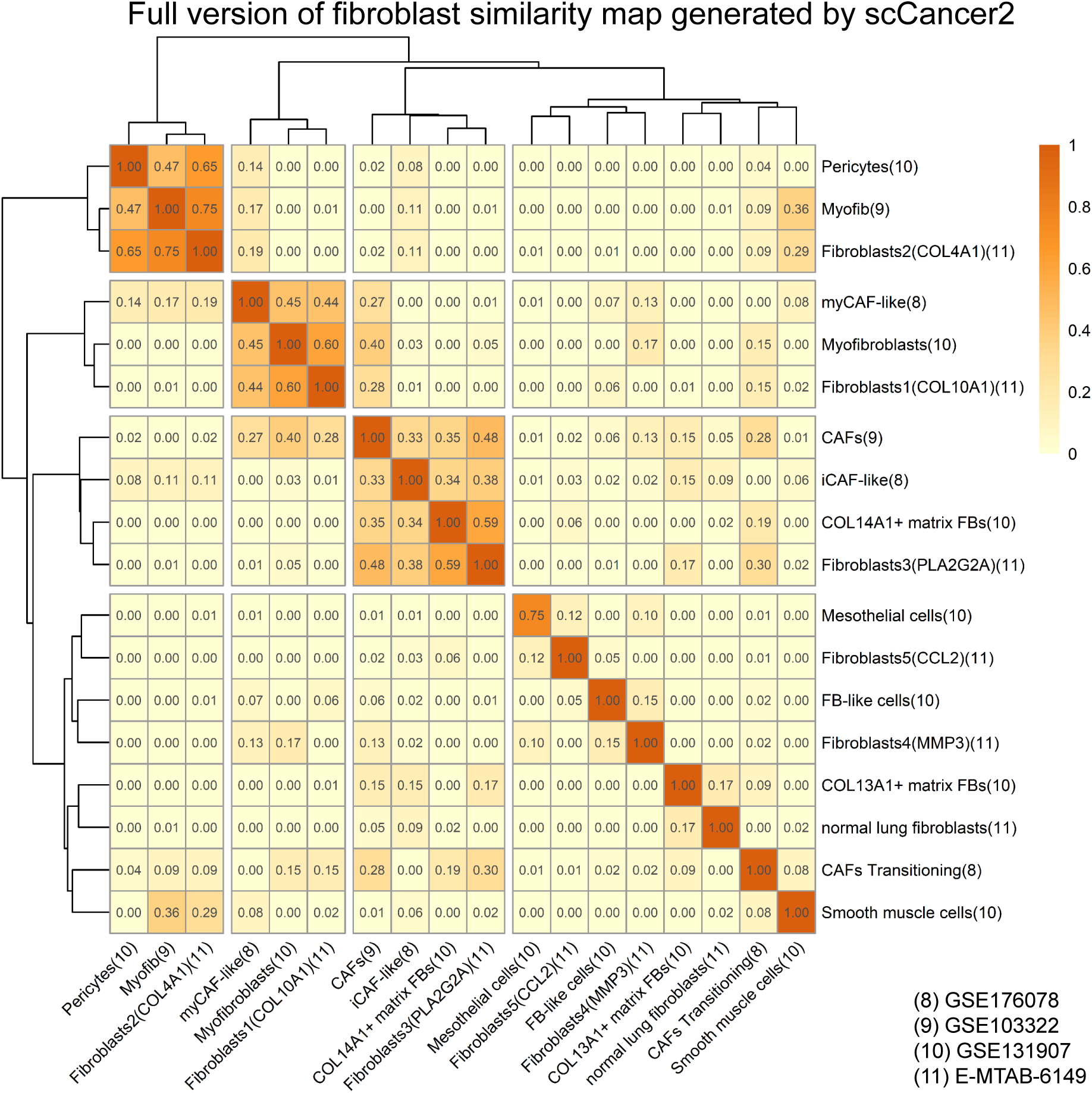
Similarity map of fibroblast subtypes generated from cross-dataset annotation (4 training sets).

**Figure S5:**
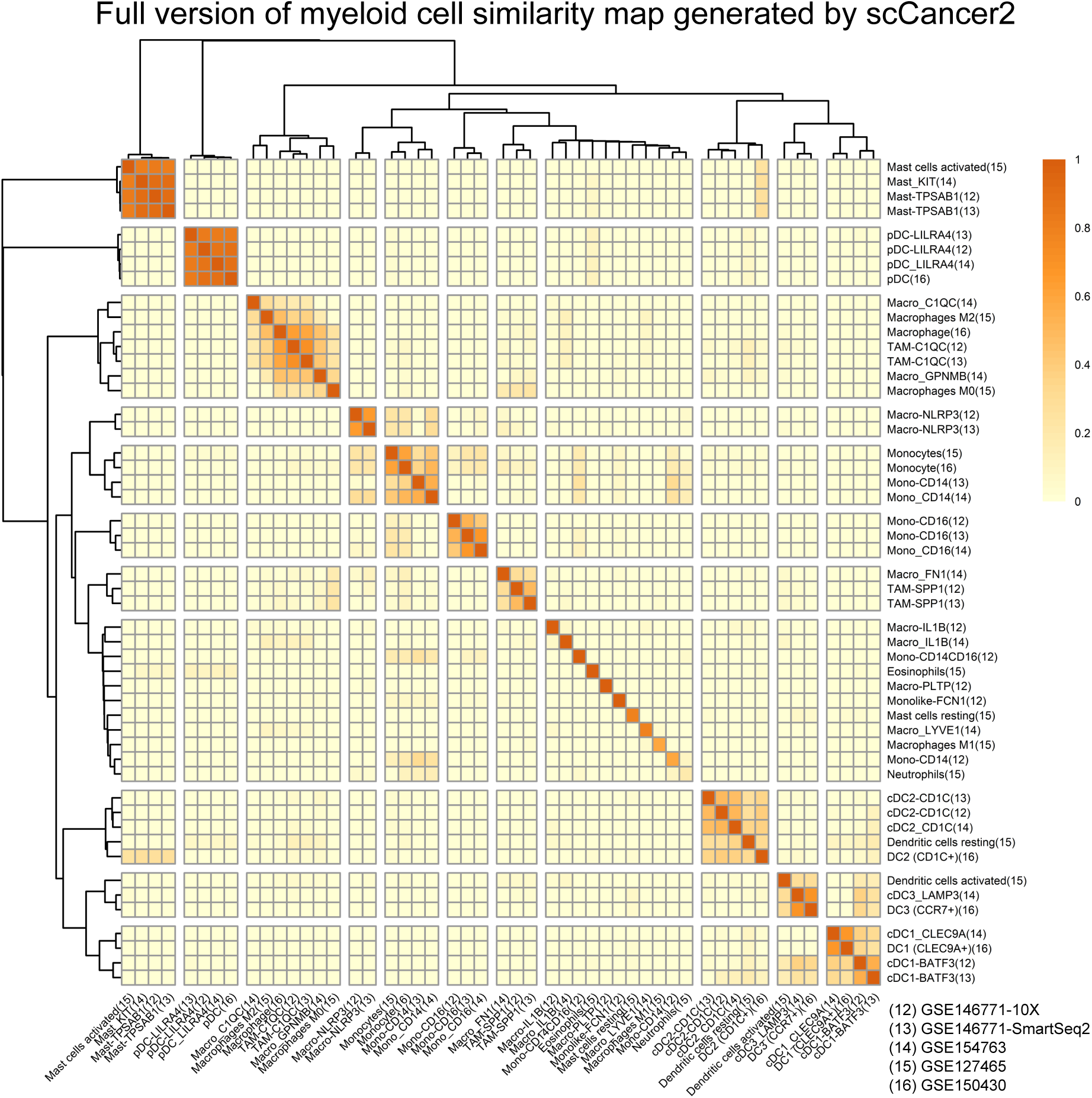
Similarity map of myeloid subtypes generated from cross-dataset annotation (5 training sets).

**Figure S6:**
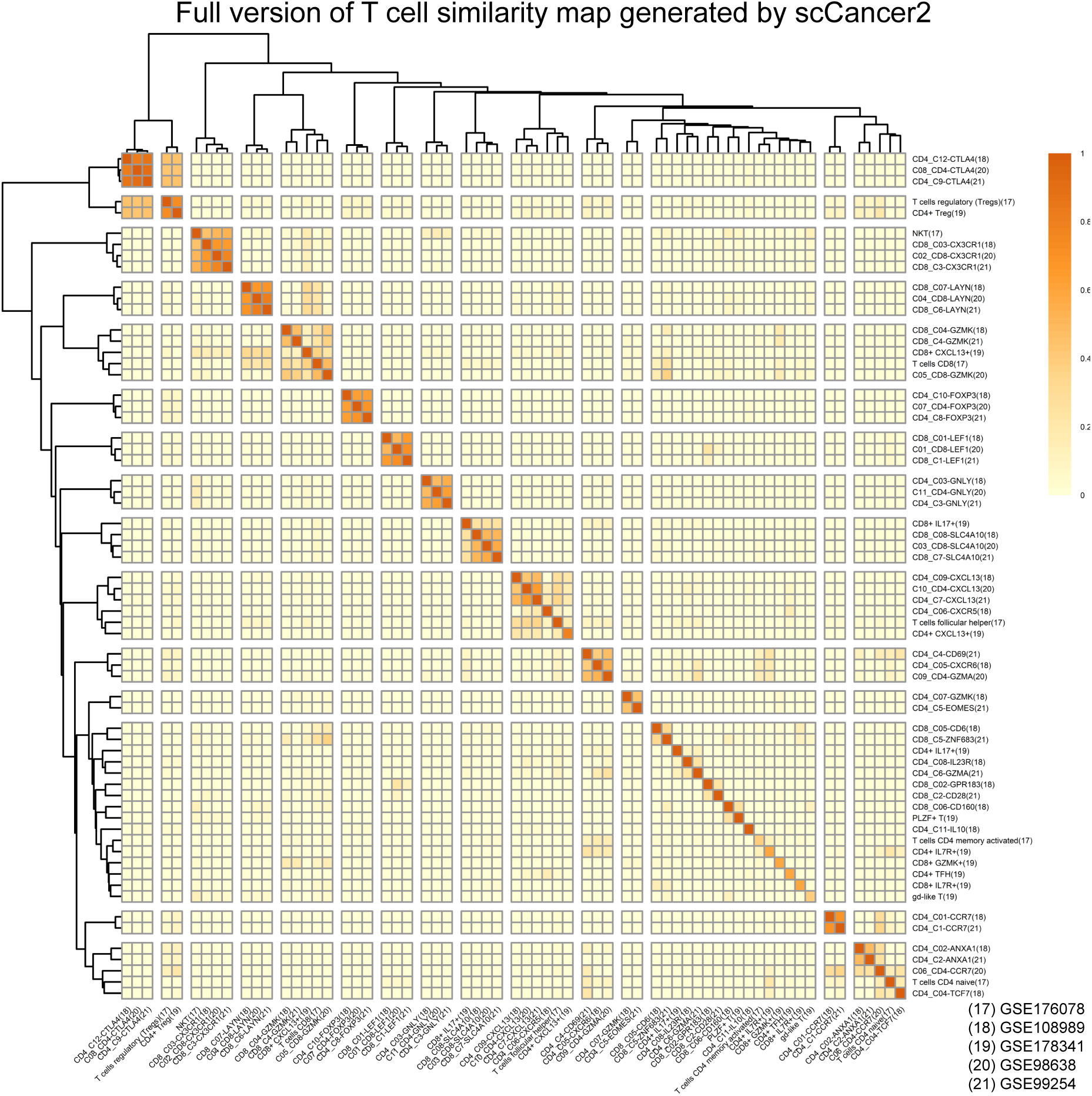
Similarity map of T cell subtypes generated from cross-dataset annotation (5 training sets).

#### More results on malignant cell identification task

**Figure S7:**
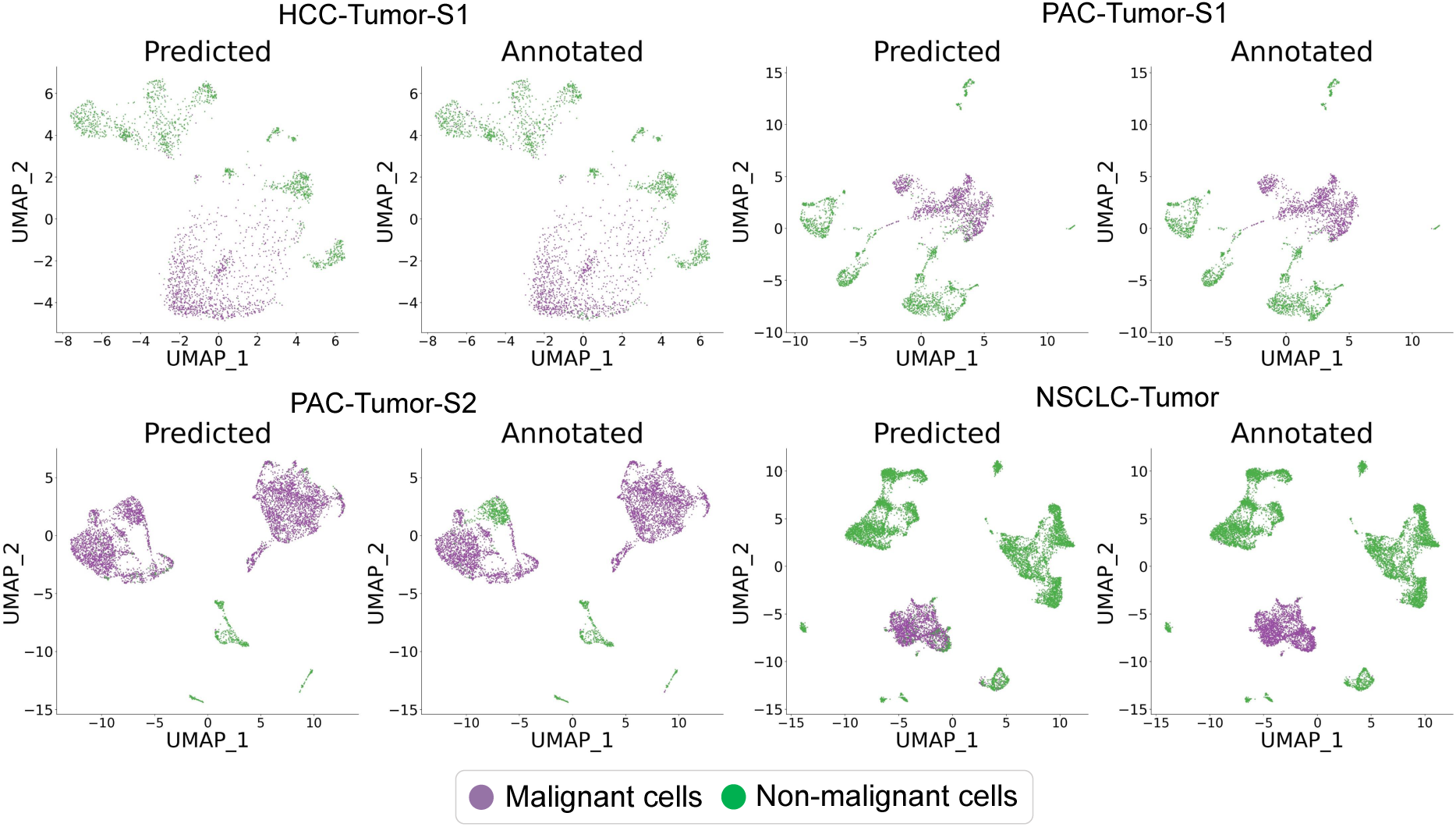
Supplemental results of case 1. Tumor samples with bimodal distribution of CNV score. The annotations of XGBoost model showed high consistency with CNV-based method.

**Figure S8:**
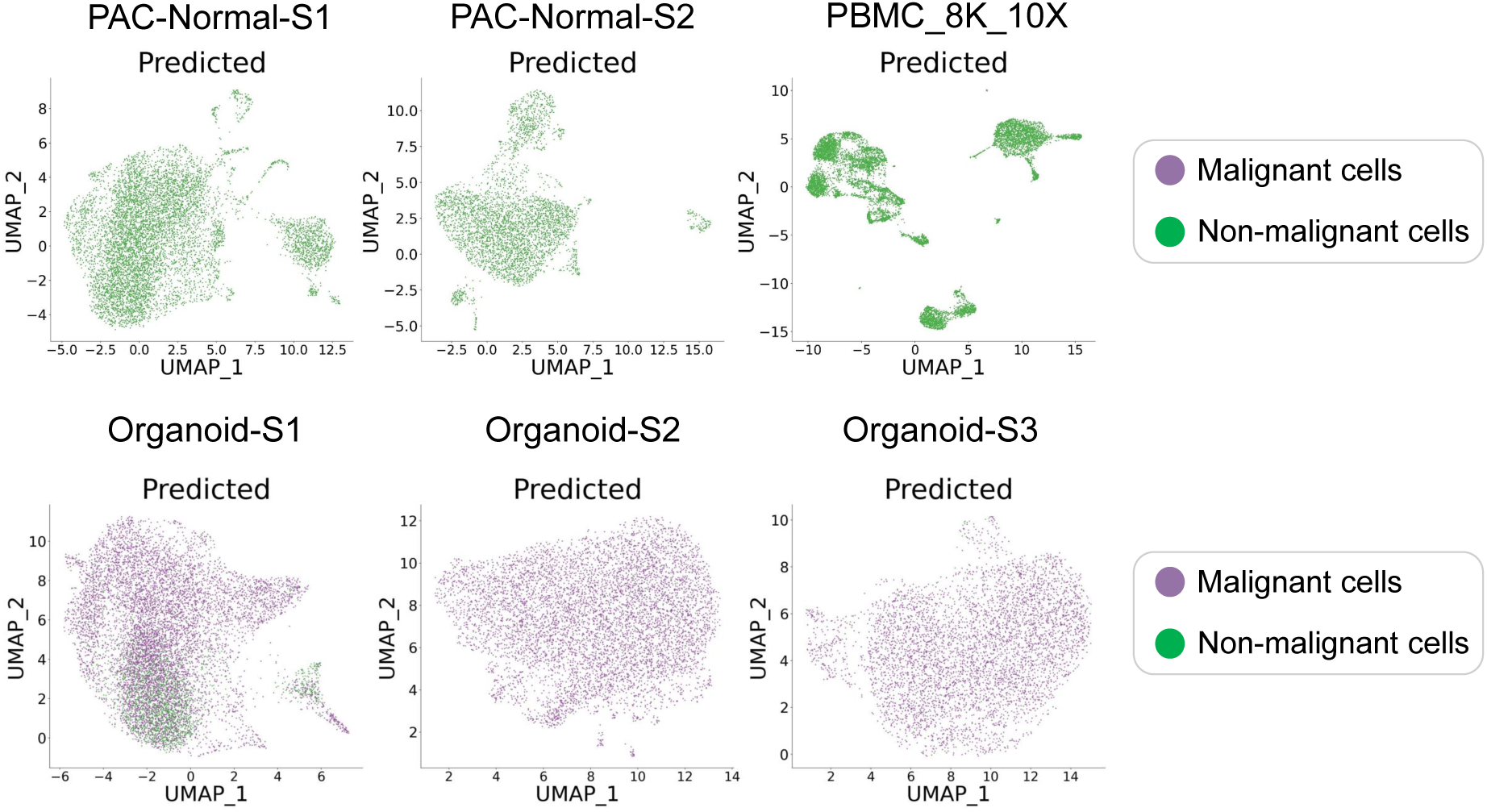
Supplemental results of case 2 and case 3. Normal samples and organoid samples. XGBoost model can effectively identify tumor or normal cells when CNV-based method fails to work.

**Figure S9:**
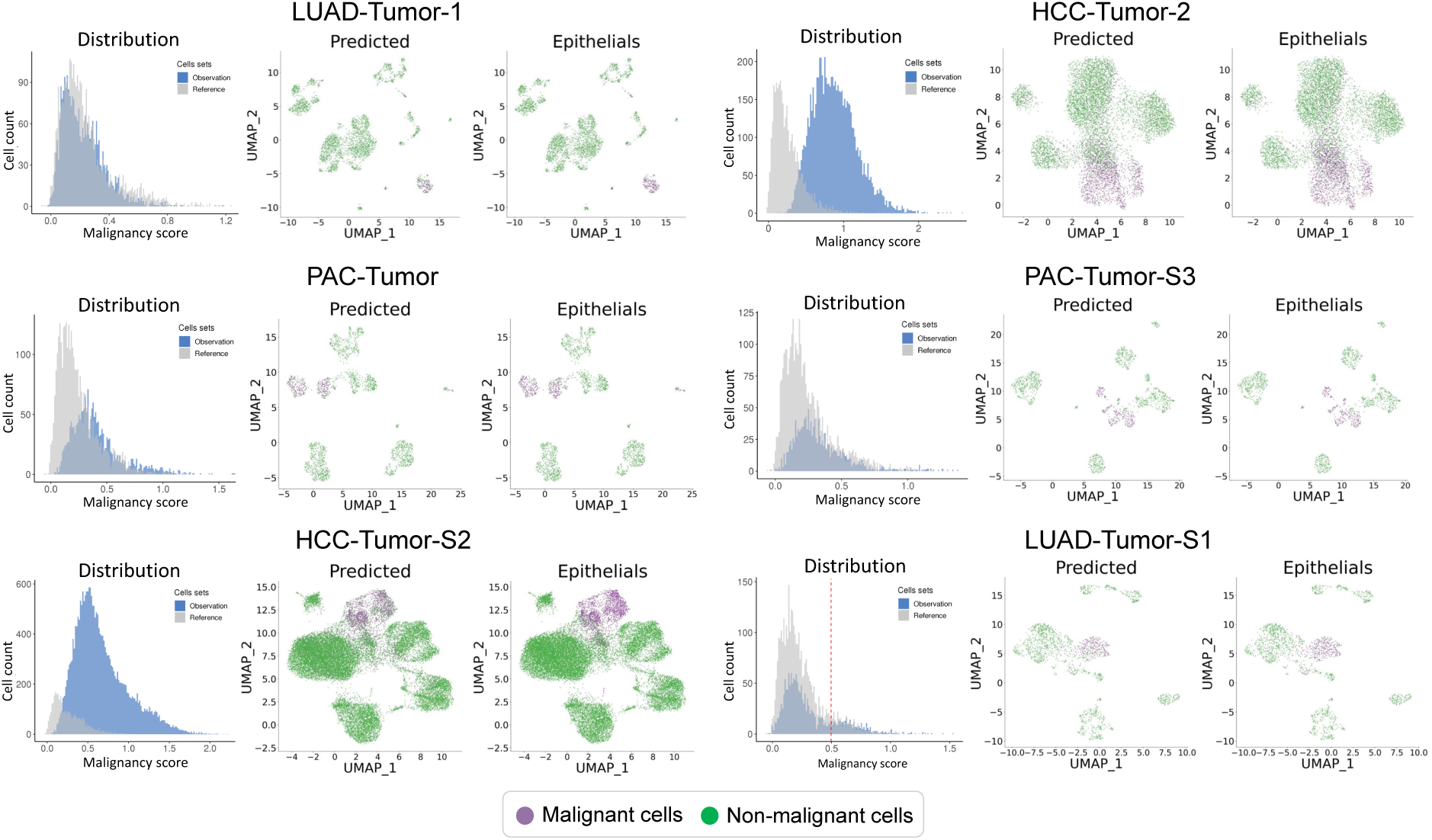
Supplemental results of case 4. Tumor samples with unimodal distribution of CNV score. XGBoost model can effectively identify malignant clusters when CNV-based method fails to work.

#### More results on spatial transcriptome analysis

**Figure S10:**
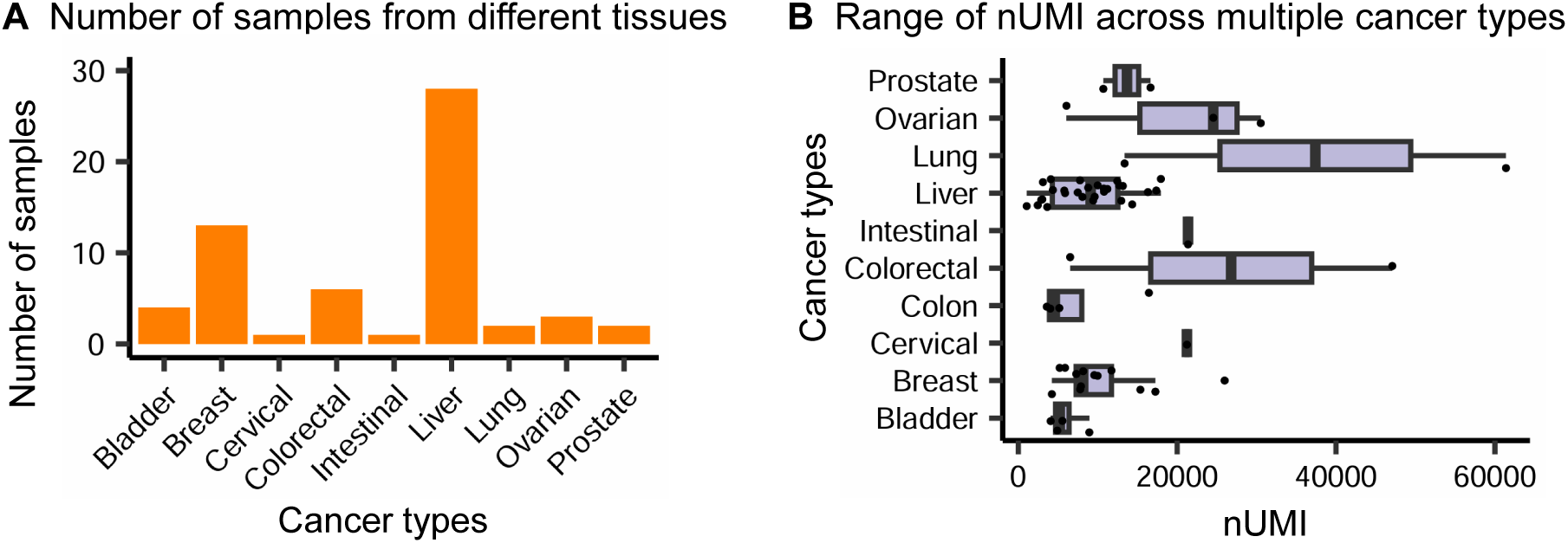
Statistical results of tumor samples from different tissues.

**Figure S11:**
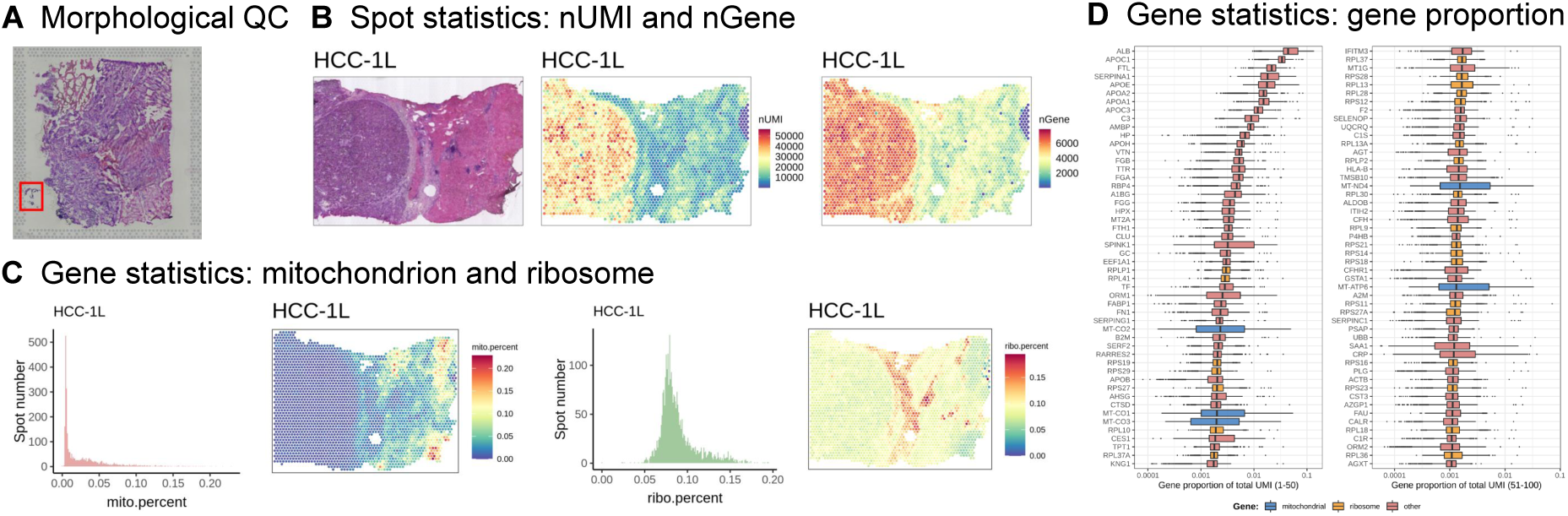
Supplemental results of stStatistics module in scCancer2 on HCC-1L.

**Figure S12:**
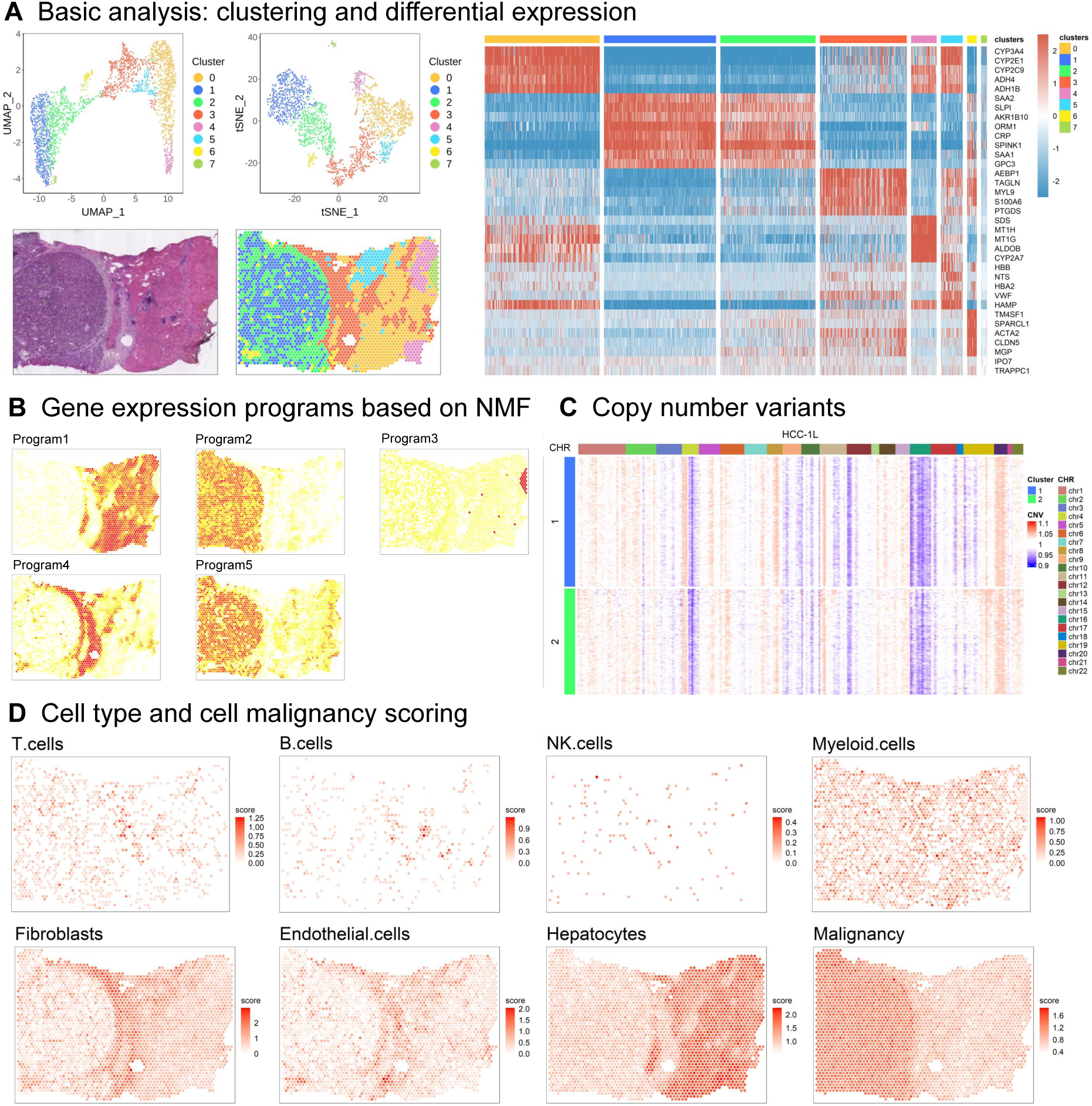
Supplemental results of stAnnotation module in scCancer2 on HCC-1L.

**Figure S13:**
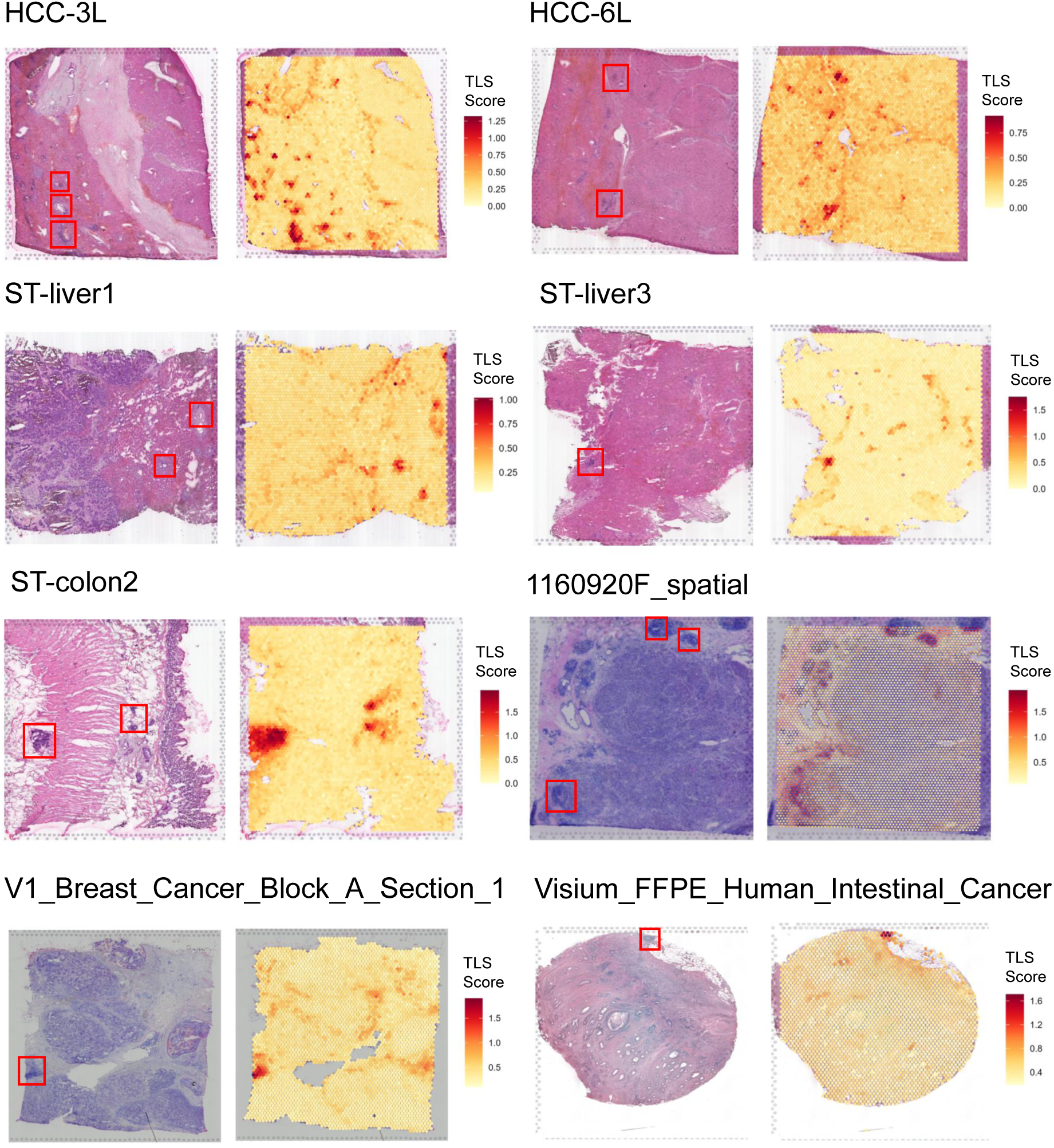
TLSs identified by scCancer2 and their corresponding H&E images. ”HCC-3L”, “HCC-6L”: Wu et al. [34]; “ST-liver1”, “ST-liver3”, “ST-colon2”: Wu et al. [40]; “1160920F spatial”: Wu et al. [31]; “V1 Breast Cancer Block A Section 1”, “Visium FFPE Human Intestinal Cancer”: 10X Genomics.

### Performance evaluation of XGBoost on cell subtype annotation

#### Performance evaluation of XGBoost on every single dataset

**Figure S14:**
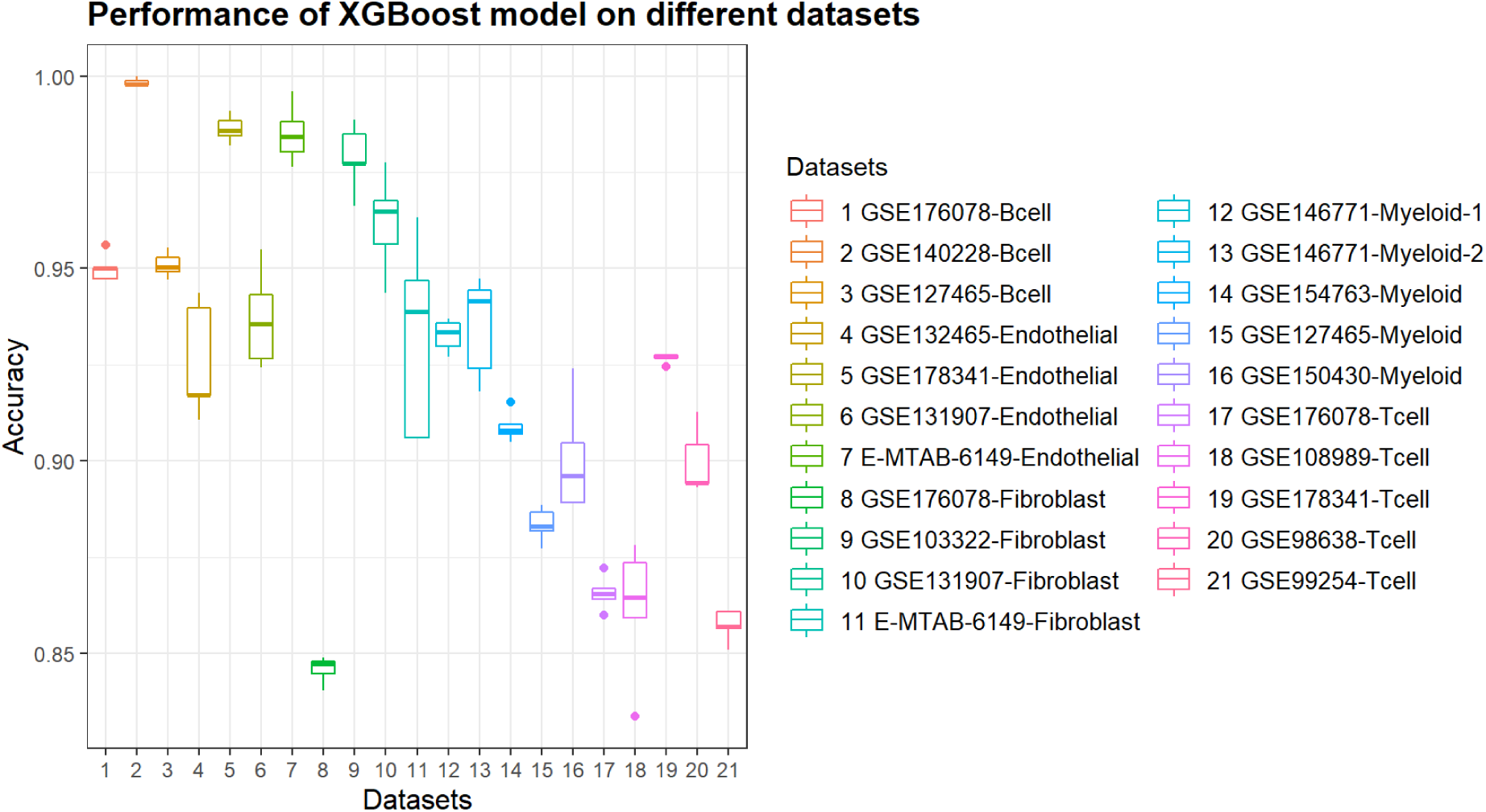
Performance evaluation of XGBoost model on subtype annotation task. The results were obtained by 5-fold cross-validation.

#### Performance evaluation of XGBoost on cross-dataset annotation

**Figure S15:**
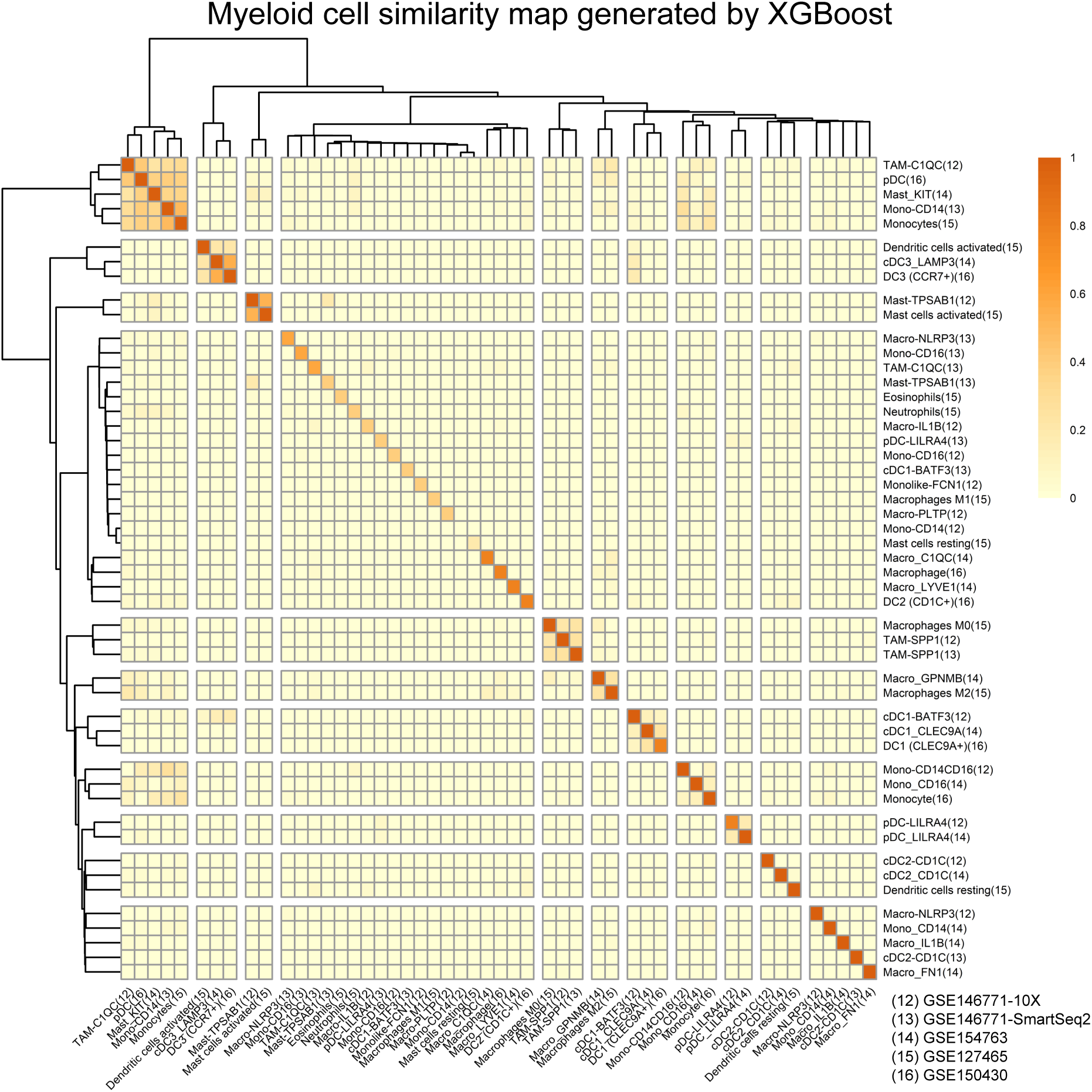
Similarity map of myeloid cell subtypes generated from cross-dataset annotation (5 training sets). XGBoost model has poor cross-dataset generalization ability on myeloid cells.

**Figure S16:**
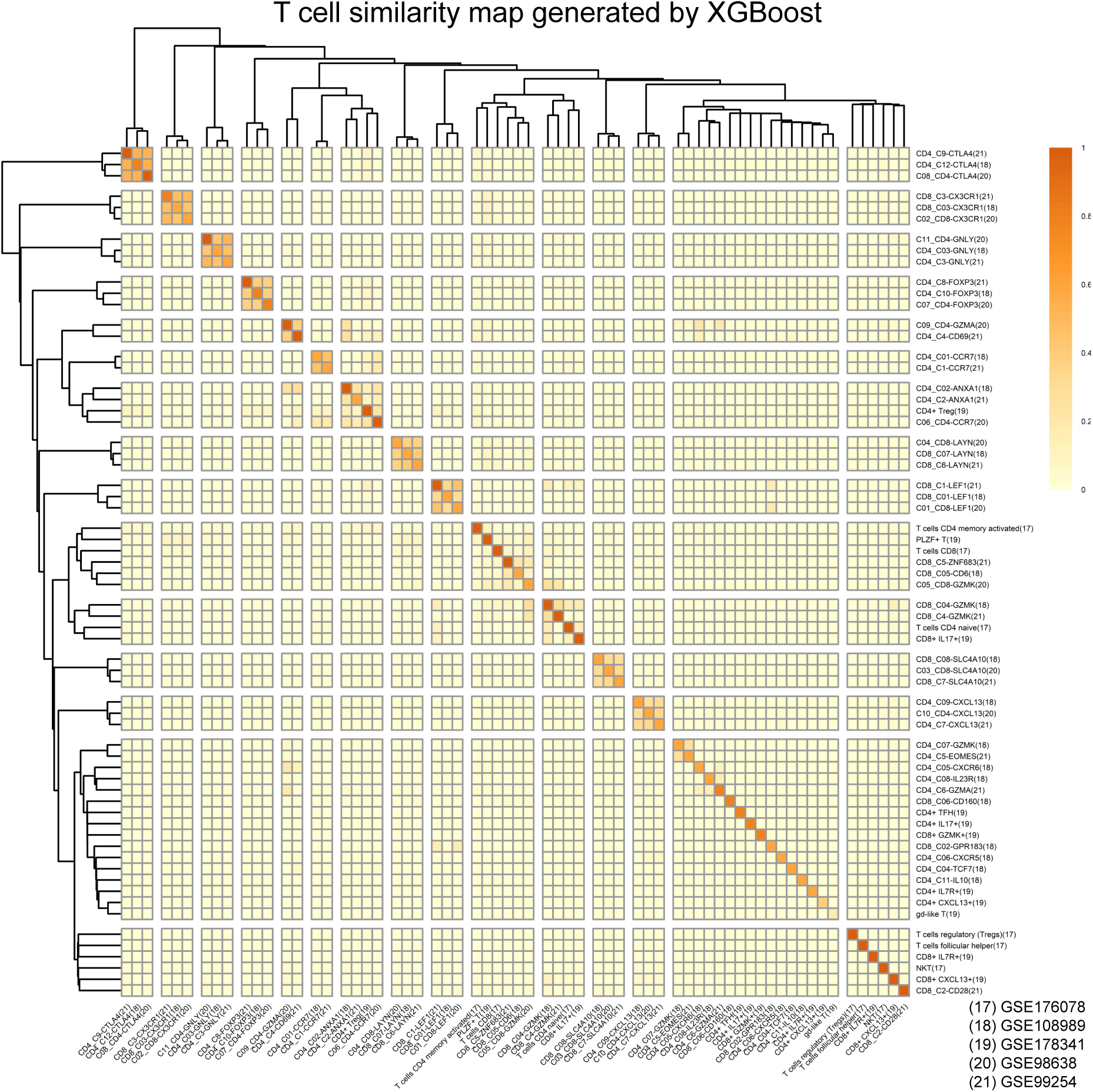
Similarity map of cell subtypes generated from cross-dataset annotation (5 training sets). XGBoost model has poor cross-dataset generalization ability on T cells.

### Supplemental Tables

**Table S1:** Methods and packages newly implemented in scCaner2.

| Steps | Packages and References | Methods |
| --- | --- | --- |
| Sample selection | Garnett [7] | Aggregated marker scores |
| Feature selection | Scibet [25] | Entropy-test |
|  | Highly Regional Genes [47] | Graph-based gene selection |
| Cell subtype annotation | Scibet [25] | Build a multinomial model for each cell subtype |
| Malignant cell identification | xgboost [12] | Extreme gradient boosting |
| Cell-cell interaction in spatial transcriptomics data | CellPhoneDB [30] | Combine the expression of multi-subunit ligand-receptor complexes |
| Copy number variation analysis | CopyKAT [11] | Calculate genomic copy number profiles of single cells and defined clonal substructure |

**Table S2:** References for cell subtype annotation in scCaner2.

| Major cell type | Accession | Reference<sup>1</sup> |
| --- | --- | --- |
| T cell (5 refs) | (1) GSE176078 | Breast2021, Wu et al. [14] |
|  | (2) GSE108989 | Colorectal2018, Zhang et al. [3] |
|  | (3) GSE178341 | Colorectal2021, Pelka et al. [13] |
|  | (4) GSE98638 | Liver2017, Zheng et al. [1] |
|  | (5) GSE99254 | Lung2018, Guo et al. [2] |
| myeloid cell (6 refs) | (1) GSE146771-10X | Colorectal2020, Zhang et al. [18] |
|  | (2) GSE146771-SmartSeq2 | Colorectal2020, Zhang et al. [18] |
|  | (3) GSE178341 | Colorectal2021, Pelka et al. [13] |
|  | (4) GSE154763 | Pan-cancer2021, Cheng et al. [15] |
|  | (5) GSE127465 | Lung2019, Zilionis et al. [16] |
|  | (6) GSE150430 | Nasopharyngeal2020, Chen et al. [17] |
| B cell (3 refs) | (1) GSE176078 | Breast2021, Wu et al. [14] |
|  | (2) GSE140228 | Liver2019, Zhang et al. [19] |
|  | (3) GSE127465 | Lung2019, Zilionis et al. [16] |
| endothelial cell (4 refs) | (1) GSE132465 | Colorectal2020, Lee et al. [21] |
|  | (2) GSE178341 | Colorectal2021, Pelka et al. [13] |
|  | (3) GSE131907 | Lung2020, Kim et al. [22] |
|  | (4) E-MTAB-6149 | Lung2018, Lambrechts et al. [23] |
| fibroblast (3 refs) | (1) GSE103322 | HeadandNeck2017, Puram et al. [20] |
|  | (2) GSE131907 | Lung2020, Kim et al. [22] |
|  | (3) E-MTAB-6149 | Lung2018, Lambrechts et al. [23] |

<sup>1</sup>For each major cell lineage, the order of references corresponds to the order of annotated labels in scCancer2.

**Table S3:**
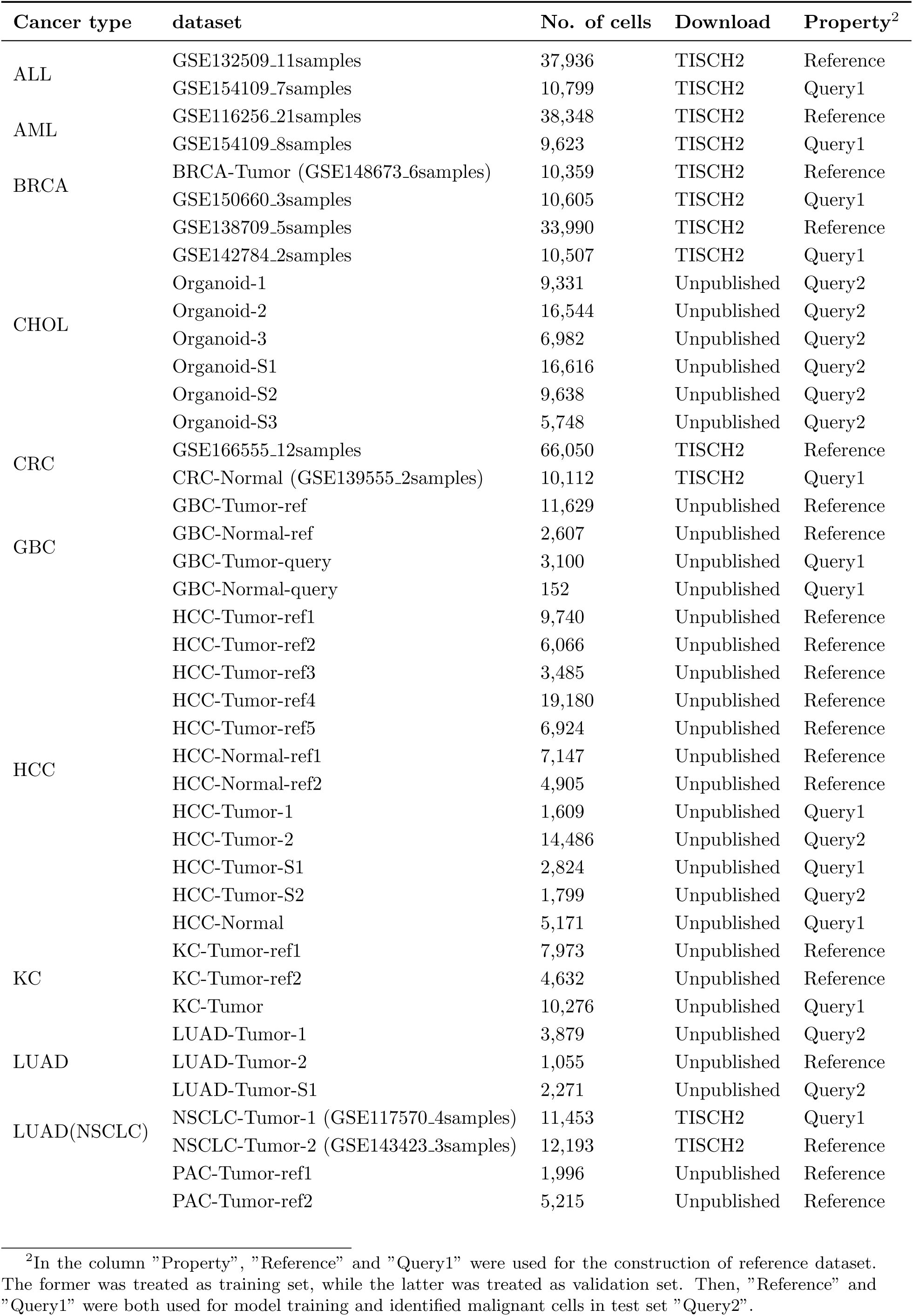
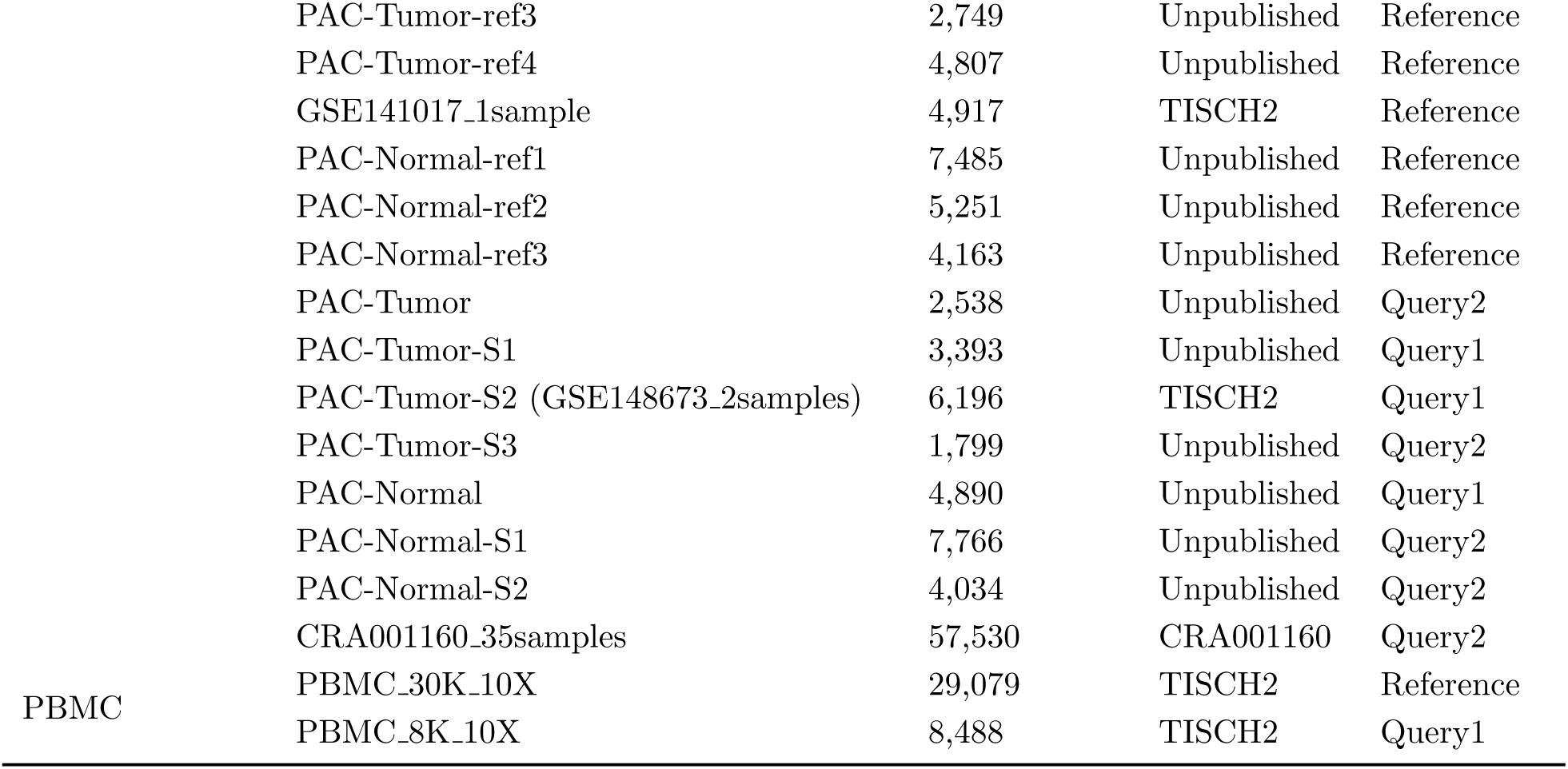
The list of scRNA-seq datasets used in malignant cell identification.

**Table S4:** The list of TCGA datasets used in malignant cell identification.

| Cancer type | No. of samples | No. of malignant samples<sup>3</sup> | No. of normal samples<sup>4</sup> |
| --- | --- | --- | --- |
| BRCA | 1222 | 1109 | 113 |
| KIRC | 611 | 539 | 72 |
| LUAD | 594 | 535 | 59 |
| LUSC | 551 | 502 | 49 |
| PRAD | 551 | 499 | 52 |
| HNSC | 546 | 502 | 44 |
| COAD | 521 | 480 | 41 |
| SKCM | 472 | 471 | 1 |
| BLCA | 433 | 414 | 19 |
| LIHC | 424 | 374 | 50 |
| STAD | 407 | 375 | 32 |
| KIRP | 321 | 289 | 32 |
| PAAD | 182 | 178 | 4 |
| READ | 177 | 167 | 10 |

<sup>3</sup>The tissue types named 'Additional New Primary', 'Additional Metastatic', 'Metastatic', 'Primary Tumor', or 'Recurrent Tumor' were labeled as "malignant".
<sup>4</sup>The tissue type named 'Solid Tissue Normal' was labeled as "nonMalignant".

**Table S5:** The list of spatial transcriptomics datasets used in scCancer2.

| Tissue types | No. of samples | Sources<sup>5</sup> |
| --- | --- | --- |
| Liver | 28 | Wu et al. [34]<sup>6</sup>, Wu et al. [40]<sup>7</sup> |
| Breast | 13 | Wu et al. [14], 10X Genomics<sup>8</sup> |
| Bladder | 4 | Gouin et al. [52] |
| Colorectal | 6 | Wu et al. [40], 10X Genomics |
| Ovarian | 3 | 10X Genomics |
| Lung | 2 | 10X Genomics |
| Prostate | 2 | 10X Genomics |
| Cervical | 1 | 10X Genomics |
| Intestinal | 1 | 10X Genomics |

<sup>5</sup>Access to 3 datasets in **Figure 5** are listed below:
<sup>6</sup>http://lifeome.net/supp/livercancer-st/data.htm
<sup>7</sup>http://www.cancerdiversity.asia/scCRLM/
<sup>8</sup>https://www.10xgenomics.com/resources/datasets/

